# Evaluation of the Efficacy and Safety of a Clinical Grade Human Induced Pluripotent Stem Cell-Derived Cardiomyocyte Patch: A Pre-Clinical Study

**DOI:** 10.1101/2021.04.07.438744

**Authors:** Shigeru Miyagawa, Takuji Kawamura, Emiko Ito, Maki Takeda, Hiroko Iseoka, Junya Yokoyama, Akima Harada, Noriko Mochizuki-Oda, Yukiko Imanishi-Ochi, Junjun Li, Masao Sasai, Fumiyo Kitaoka, Masaki Nomura, Naoki Amano, Tomoko Takahashi, Hiromi Dohi, Eiichi Morii, Yoshiki Sawa

**Author notes:** Corresponding Author: Yoshiki Sawa, M.D., Ph.D., Department of Cardiovascular Surgery, Osaka University Graduate School of Medicine 2-2 Yamadaoka, Suita, Osaka 565-0871, Japan.

## Abstract

**Aims:** Cardiomyocyte-derived induced pluripotent stem cells (iPSCs) may represent a promising therapeutic strategy for severely damaged myocardium. This study aimed to assess the efficacy and safety of clinical grade human iPSC-derived cardiomyocyte (hiPSC-CM) patches and conduct a pre-clinical proof-of-concept analysis.

**Methods and results:** A clinical grade hiPSC line was established from peripheral blood mononuclear cells collected from a healthy volunteer homozygous for human leukocyte antigens and differentiated into cardiomyocytes using cytokines and chemical compounds. hiPSC-CMs were cultured on temperature-responsive culture dishes to fabricate the hiPSC-CM patch. The hiPSC-CMs expressed cardiomyocyte-specific genes and proteins while electrophysiological analyses revealed that hiPSC-CMs were similar to the human myocardium. *In vitro* safety studies using cell growth, soft agar colony formation, and undifferentiated cell assays indicated that tumourigenic cells were not present. Moreover, no genomic mutations were discovered using whole genome and exome sequencing analysis. Tumour formation was not detected in an *in vivo* tumourigenicity assay using NOG mice. General toxicity tests also showed no adverse events due to hiPSC-CM patch transplantation. An efficacy study using a porcine model of myocardial infarction demonstrated significantly improved cardiac function with angiogenesis and a reduction in interstitial fibrosis, which was enhanced by cytokine secretion from hiPSC-CM patches after transplantation. No lethal arrhythmias were observed.

**Conclusion:** hiPSC-CM patches show promise for future translational research and clinical trials for ischaemic heart failure.

**One-sentence summary:** This pre-clinical study provides a proof-of-concept of the safety and efficacy of hiPSC-CM patches for the treatment of heart failure.

**Translational Perspective:** Regenerative therapy using cells and tissues is attractive as a novel approach for treating severe heart failure. We focused on human iPS cell-derived cardiomyocytes (hiPSC-CMs) as a cell source. Using basic research, the characteristics of hiPSC, hiPSC-CMs, and hiPSC-CM patches were determined *in vitro* and *in vivo*. We also conducted a pre-clinical study using a porcine model of myocardial infarction that confirmed the safety and efficacy of the hiPSC-CM patch, highlighting its potential for clinical application.

## Introduction

In spite of advances in medical treatment, the fatality rate of heart failure remains high. The treatment of heart failure would benefit from novel clinical equipment and techniques. In recent years, research on human induced pluripotent stem cells (hiPSCs) has shown that they may be a promising source of stem cells that can replace lost cells in diseased organs^1–3^. Moreover, basic research has proven that hiPSCs can also supply cardiomyocytes to the failed heart through myocardial tissue such as a cardiomyocyte patch, indicating that these cells may be introduced as clinical treatment for heart failure^4–7^. On the other hand, there are major concerns regarding the safety of hiPSCs in clinical applications, especially concerning tumourigenicity ^8–10^. Adequate pre-clinical study concerning the safety and efficacy *in vitro* and *in vivo* may show the promise of good clinical results for the eventual clinical trials of this technique for heart failure patients.

In this study, we aimed to demonstrate whether a clinical grade hiPSC-derived cardiomyocyte (hiPSC-CM) patch can act as functional myocardial tissue. We performed a pre-clinical study to ensure safety and conducted a proof-of-concept analysis of hiPSC-CM patches in clinical applications.

## Methods

### Clinical grade hiPSCs

A clinical grade hiPSC line (QHJI14s04) was established from the peripheral blood mononuclear cells of a healthy volunteer donor homozygous for human leukocyte antigen (HLA) and with the most frequent haplotype in the Japanese population^11^ in the cell processing centre of the Center for iPS Cell Research and Application (Kyoto University, Kyoto, Japan). We obtained informed consent from the donor and all studies complies with the Declaration of Helsinki. To ensure cell quality, we performed characterisation tests as described in the online Supplementary material.

### Generation of the master cell bank (MCB)

We received two cryovials of hiPSC passage (P) 10 from CiRA. One vial was used for evaluation of the growth rate and optimisation of the culture conditions. The other cryovial was used for producing the MCB. The MCB was prepared from cryopreserved P12 cells using qualified reagents and raw materials in the Center for Gene and Cell Processing of Takara Bio Inc. (Kusatsu, Japan), which complies with good manufacturing practice (GMP)/GCTP standards. The cells and supernatants used for analysis were prepared by sub-culturing P12 cells for two additional passages. In accordance with ICH Q5A and 5D, the MCB was inspected by DNA fingerprinting and electron microscopy and evaluated for sterility, reverse transcriptase activity, and the presence of mycoplasma, human viruses, or infectious retroviruses (*in vitro* and *in vivo*).

### Cardiomyogenic differentiation, purification, and elimination of residual undifferentiated hiPSCs

The hiPSC line QHJI14s04 was cultured on iMatrix511 (Nippi, Tokyo, Japan)-coated dishes in Stem Fit Ak03N (Ajinomoto, Tokyo, Japan), followed by cardiac differentiation, purification, and elimination of residual undifferentiated hiPSCs. The cells were also frozen. Methods are detailed in the online Supplementary material.

### hiPSC-CM patch preparation

Prior to cell seeding, the surfaces of temperature-responsive dishes (UpCell; CellSeed, Tokyo, Japan) were coated with Dulbecco’s modified Eagle’s medium (DMEM; Nacalai Tesque, Kyoto, Japan) supplemented with 20% foetal bovine serum (FBS; Sigma-Aldrich, St. Louis, MO) overnight. After freeze-thawing, the hiPSC-CMs were plated onto the UpCell dishes in DMEM containing 20% FBS and cultured at 37 °C and 5% CO_2_. After 72 h in culture, the hiPSC-CM patches were harvested and washed gently with Hanks’ balanced salt solution (+).

### Flow cytometry

Cells were stained with antibodies for cTNT, αSMA, vimentin, and CD31, incubated with fluorescently conjugated secondary antibodies, and assessed using the FACS Canto II system (BD Biosciences, Franklin Lakes, NJ). Method is detailed in the online Supplementary material.

### RNA isolation and quantitative polymerase chain reaction (qPCR)

Total RNA was extracted from the cells and heart tissues, and then reverse-transcribed into cDNA. qPCR was performed with the ViiA 7™ Real-Time PCR or ABI PRISM 7700 systems (Applied Biosystems, Foster City, CA). Methods are detailed in the online Supplementary material.

### PCR arrays

Total RNA was extracted from hiPSCs and hiPSC-CMs. Human adult and foetal heart total RNA was purchased from Takara Bio. For the analysis of human stem cell-associated genes, the ViiA 7™ Real-Time PCR system was used to run the Human Stem Cell RT² Profiler™ PCR Array (Qiagen, Hilden, Germany). For the analysis of cardiac differentiation-associated genes, gene expression was analysed using the TaqMan™ Array Human Cardiomyocyte Differentiation by BMP Receptors (Thermo Fisher Scientific, Waltham, MA). Methods are detailed in the online Supplementary material.

### Single-cell RNA sequencing

The single-cell sequencing library was generated using the ICELL8 cx platform (Takara Bio) and sequencing was performed using the HiSeq 3000 sequencer (Illumina, San Diego, CA). Methods are detailed in the online Supplementary material.

### Whole genome/exome sequencing analysis

Whole genome sequencing (WGS), whole exome sequencing (WES), and single nucleotide polymorphism (SNP) array experiments were performed using peripheral blood mononuclear cells from the donor (termed origin or control), the MCB, expansion cultures of the MCB, hiPSC-CMs, and hiPSC-CM patches. WGS libraries were generated with the KAPA Hyper Prep Kit (Kapa Biosystems, Wilmington, MA) from fragmented genomic DNA sheared by Covaris LE220 (Covaris, Brighton, UK). For WES, adapter-ligated libraries were prepared with the KAPA Hyper Prep Kit and sequencing libraries were constructed using SeqCap EZ Human Exome Library v3.0 (Roche, Basel, Switzerland). Cluster generation was performed with the HiSeq PE Cluster Kit v4 (Illumina) using Illumina cBot. Sequencing was performed using the HiSeq2500 platform (Illumina) in the 126 paired-end mode. Methods are detailed in the online Supplementary material.

### SNP array analysis

Copy number variations (CNVs) were called using the HumanOmniExpress24 v1.1 genotyping array (Illumina). We prepared 200 ng of genomic DNA, which was hybridised using the HumanOmniExpress24 v1.1 DNA Analysis Kit (Illumina), and evaluated its intensity using the iScan (Illumina). After exporting a final report using GenomeStudio (v2011.1; Illumina), CNV analyses were performed using PennCNV (v1.0.3)^12^, Mosaic Alteration Detection-MAD (v1.0.1) ^13^, and GWAS tools (v1.16.1) ^14^; the test samples were compared with the control sample and the called CNVs were curated manually.

### Measurement of cytokine levels

The supernatants of cell patches were collected after culturing under normoxic or hypoxic (5% O_2_) conditions and analysed using a fluorescence-dyed microsphere-based immune assay (Bio-Rad Laboratories, Hercules, CA) following the manufacturer’s instructions. The concentration of cytokines, such as angiogenin, angiopoietin-1, angiopoietin-2, hepatocyte growth factor (HGF), and vascular endothelial growth factor (VEGF), was measured and analysed using the Bio-plex suspension array system (Bio-Rad Laboratories).

### Transmission electron microscopy

The samples were immersed in 0.5% uranyl acetate (Fujifilm Wako Pure Chemical Corporation, Osaka, Japan), dehydrated in ethanol (Muto Pure Chemicals, Tokyo, Japan) and propylene oxide (Sigma-Aldrich), and then embedded in epoxy resin. Ultrathin sections were cut using an EM UC7 ultra-microtome (Leica Microsystems, Wetzlar, Germany). The sections were imaged using an H-7500 transmission electron microscope (Hitachi High-Technologies, Tokyo, Japan). Methods are detailed in the online Supplementary material.

### Electrophysiological properties of the hiPSC-CM patches

The hiPSC-CM patches were transferred onto a MED probe (Alpha MED Scientific, Osaka, Japan) and incubated until they were attached. Extracellular field potentials were monitored using a multi-electrode array (MEA) system (MED64; Alpha MED Scientific), recorded for 10 min, and analysed using Mobius software (Alpha MED Scientific).

### Mechanical properties of the hiPSC-CM patches

Mechanical properties were assessed using the MicroTester G2 (CellScale, Waterloo, ON, Canada). hiPSC-CM patches were transferred into a heated bath containing culture medium at 37 °C. The two ends of each hiPSC-CM patch were hooked to a fixed wire and a force-sensing probe. The force generated by the patch was calculated based on the displacement of the probe. The probe was moved downward to stretch the patch to different extents until the tissue broke. The force data were recorded at a sample rate of 5 Hz.

### Intracellular calcium activity and contraction properties of hiPSC-CMs

hiPSC-CMs were seeded onto 96-well plates at 1–2 × 105 cells/well. Intracellular calcium activity was analysed with FDSS/μCELL (Hamamatsu Photonics, Shizuoka, Japan) and contraction properties were monitored with a Cell Motion Imaging System (SI8000; Sony Biotechnology, Tokyo, Japan). Methods are detailed in the online Supplementary material.

### Generation of the porcine chronic myocardial infarction model and transplantation of hiPSC-CM patches

All experimental procedures were performed according to the national regulations and guidelines, reviewed by the Committee for Animal Experiments, and finally approved by the president of Osaka University. A chronic myocardial infarction model was constructed in mini-pigs. Animals were randomly divided into hiPSC-CM patch transplantation group or a sham operation group. Cardiac function was evaluated by echocardiography, selective coronary angiography, cardiac magnetic resonance imaging (MRI), and telemetered Holter electrocardiography. Methods, including experimental animal model generation, cardiac function assessment, and histological, molecular biology, and statistical analyses are detailed in the online Supplementary material.

### Cell growth assay

hiPSC-CMs were cultured in DMEM supplemented with 10% FBS at 37 °C and 5% CO_2_. After reaching 80–90% confluency, the cells were harvested using 0.05% trypsin-EDTA (Thermo Fisher Scientific) and passaged five times; the growth rate (R*_n_*) at passage = *n*, i.e. doubling per day, was calculated using the following formula:

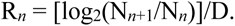

where N_n_ is the harvested cell number at passage *n* and D is the culture day.

### Soft agar colony formation assay

The soft agar colony formation assay was conducted to rule out tumourigenicity. Methods are detailed in the online Supplementary material.

### General toxicity tests and tumourigenicity assay

The hiPS-CM patches were tested for general toxicity and tumourigenicity using immunodeficient NOD/Shi-scid, IL-2R γnull mice (NOG mice; In-Vivo Science Inc., Kawasaki, Japan). For general toxicity testing, hiPSC-CM patches consisting of 1.9 × 10^6^ hiPSC-CMs were directly transplanted onto the surface of the left anterior wall of the heart. Mice were euthanised and dissected 28 days after transplantation. Gross abnormalities and the weights of the major organs were recorded. Haematological and biochemical evaluations were conducted using peripheral blood. For the tumourigenicity assay, hiPSC-CM patches consisting of 1.9 × 10^6^ hiPSC-CMs with or without purification and elimination of residual undifferentiated hiPSCs were directly transplanted onto the surface of the left anterior wall of the heart. Mice were euthanised and dissected 16 weeks after transplantation. The major organs and tissues were carefully inspected, and any gross pathological findings were collected and stored for further examination. Staining protocols are detailed in the online Supplementary material.

### Histological analysis

All autopsy tissue specimens of murine and porcine hearts transplanted with hiPSC-CM patches were fixed in 10% buffered formalin (Fujifilm) and embedded in paraffin using a Microm STP 120 Spin Tissue Processor (STP120-3; Thermo Fisher Scientific). Picrosirius red staining (Fujifilm) was performed on serial paraffin-embedded sections. Haematoxylin and eosin staining was performed along with immunostaining using anti-Ki-67 and anti-lamin antibodies (Table S1). The sections were assessed under a light microscope (Leica). Staining protocols are detailed in the online Supplementary material.

### Immunofluorescent staining

Cardiomyocyte aggregates, hiPSC-CM patches, and excised heart samples were fixed in 4% paraformaldehyde, frozen in liquid nitrogen, and cryosectioned. Immunofluorescent staining was performed with the primary and secondary antibodies listed in Table S1. Cell nuclei were counterstained with Hoechst 33342 (1:100; Dojindo, Kumamoto, Japan) and the sections imaged using a confocal laser scanning microscope (FV10i; Olympus, Tokyo, Japan). The system was controlled with FV10-ASW 3.1 software (Olympus).

### Statistical analysis

Statistical significance of *in vitro* experiments was determined by a two-tailed Student’s *t*-test. JMP Pro 13 software (SAS Institute Inc., Cary, NC) was used for statistical analysis of the *in vivo* efficacy trial of porcine species. Continuous values are expressed as the mean ± standard deviation (SD). The analyses were performed using nonparametric methods because sample sizes were too small to determine a normal or skewed distribution. Within-group differences were compared using the Wilcoxon signed-rank test; between-group differences were compared with the Wilcoxon-Mann-Whitney U test. *P* values < 0.05 were considered statistically significant.

## Results

### Establishing the MCB

The human iPS cell line QHJI14s04 was established as clinical grade hiPSCs by transferring multiple genes into peripheral blood mononuclear cells collected from a healthy volunteer homozygous for HLA (Table S2). In the process of establishing hiPSCs and MCB, quality checks were performed at each point (Figure S1). The quality of hiPSCs was assessed by determining colony morphology; the residual plasmid vector used for iPS production; karyotype; disappearance of the expression of pluripotent markers such as *POU5F1*, *NANOG*, *TRA-1–60*, *TRA-2–49*, and *SSEA-4*; sterility, absence of mycoplasma, endotoxins, and viruses; HLA typing, and short tandem repeat (STR) genotyping (Table S3). We generated the MCB using culture-expanded hiPSCs produced under GMP conditions. The quality of the MCB was confirmed as there was no contamination by foreign pollutants such as bacteria, mycoplasma, or viruses, and the MCB was further analysed via STR genotyping (Table S4).

### Characterisation of hiPS-CMs and hiPS-CM patches

In the process of producing hiPSC-CM, and hiPSC-CM patch quality checks were performed at each point (Figure S1). The quality of the hiPSC-CM was confirmed by cell viability, purity of cardiomyocytes, sterility, and absence of mycoplasma and endotoxins (Table S5). During myocardial differentiation, the number of cells increased. qPCR and immunohistochemistry demonstrated that the expression of markers for pluripotency, early mesoderm, cardiac progenitor cells, and cardiomyocytes changed over time (Figure S2). Analysis of undifferentiated stem cell-related genes revealed that their expression in hiPSC-CMs was lower than in hiPSCs (Figure S3A). Moreover, analysis of cardiomyocyte differentiation-related genes revealed that their expression in hiPSC-CMs was more similar to that in the human adult or foetal heart than in undifferentiated hiPSCs (Figure S3B).

hiPSC-CMs were 60–80 % positive for the cardiac troponin T (cTNT) marker. Most TNT-negative non-cardiomyocytes expressed the smooth muscle cell marker αSMA or the fibroblast marker vimentin; however, few expressed the endothelial cell marker CD31, whose positive cell rate was 1–5% (Figure 1A). Single-cell RNA-seq identified four cell populations, three of which (clusters 0, 1, and 2) consisted of cardiomyocytes that expressed the cardiac cell marker *TNNT2* and differed in their expression level of *ACTN2*. The population (cluster 3) expressed *POSTN* and *ACTA2*, which are highly expressed in fibroblasts and smooth muscle, respectively (Figure 1B). In addition, immunohistochemistry of hiPSC-CMs showed that they express ventricular muscle contractile proteins, such as myosin light chain 2v (MLC2v), beta cardiac myosin heavy chain (β-MHC), gap junction protein, and connexin 43 (Figure 1C). Furthermore, the expression of cardiomyocyte ion channels in hiPSC-CMs was similar to that of adult hearts (Figure S3C). The drug response of hiPSC-CMs was assessed by measuring calcium transients and contractile properties, and the addition of isoproterenol showed a marked positive inotropic effect. Moreover, the addition of proarrhythmic E-4031 showed a clear QT prolongation (Figure S4, S5).

**Figure 1.**
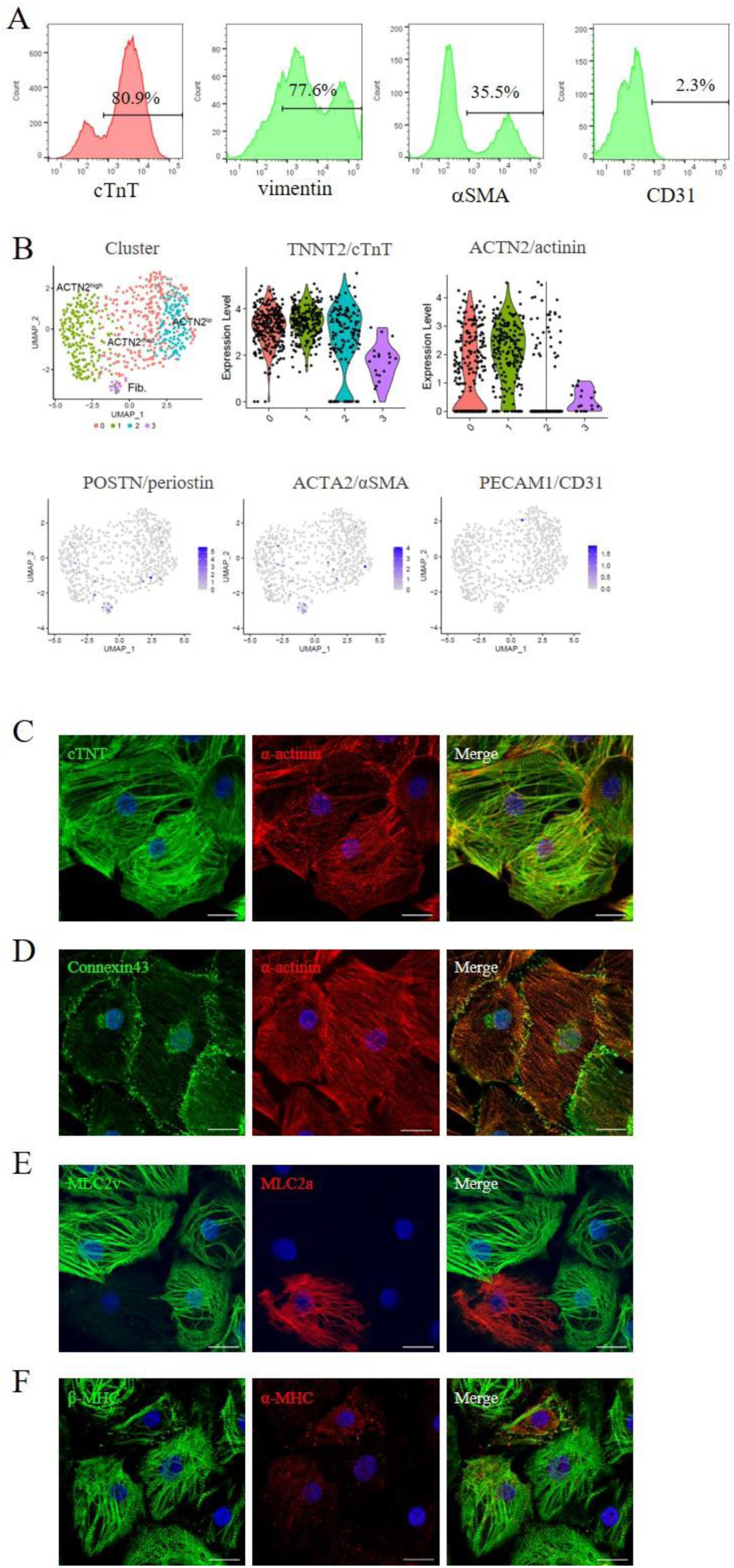
**Characterisation of hiPSC-CMs.** A: The populations within hiPSC-CMs were assessed via flow cytometry. The histograms of vimentin, αSMA, and CD31 show the data in the cTNT-negative cell population. B: Four cell populations determined by single-cell RNA-seq. C–F: The structure and morphology of hiPSC-CM. Immunofluorescence of cardiac-specific proteins: (C) cTNT (green) and α-actinin (red), (D) connexin-43 (green) and α-actinin (red), (E) MLC2v (green) and MLC2a (red), and (F) β-MHC (green) and α-MHC (red). Scale bar: 20 μm.

Prior to transplantation surgery, hiPSC-CM patches were prepared using temperature-responsive culture dishes (Figure 2A, B). Immunohistochemistry of the patches revealed well organised sarcomeric structures and the expression of extracellular matrix proteins (Figure 2C, D). Furthermore, the ultrastructure of hiPSC-CM patch showed myofibrils with transverse Z-bands and a mitochondrial structure (Figure S6A). The hiPSC-CM patch also expressed various cytokines, such as HGF and stromal cell-derived factor (SDF), and was responsive to hypoxic stimulation (Figure 3A, S6B). In addition, the hiPSC-CM patch showed synchronous, regular, and continuous beating, which indicates electrical linkage throughout the patches (Figure 3B). The hiPSC-CM patch responded according to the Frank-Starling mechanism, where the contraction force of the hiPSC-CM patch increased as its stretch rate increased (Figure 3C).

**Figure 2.**
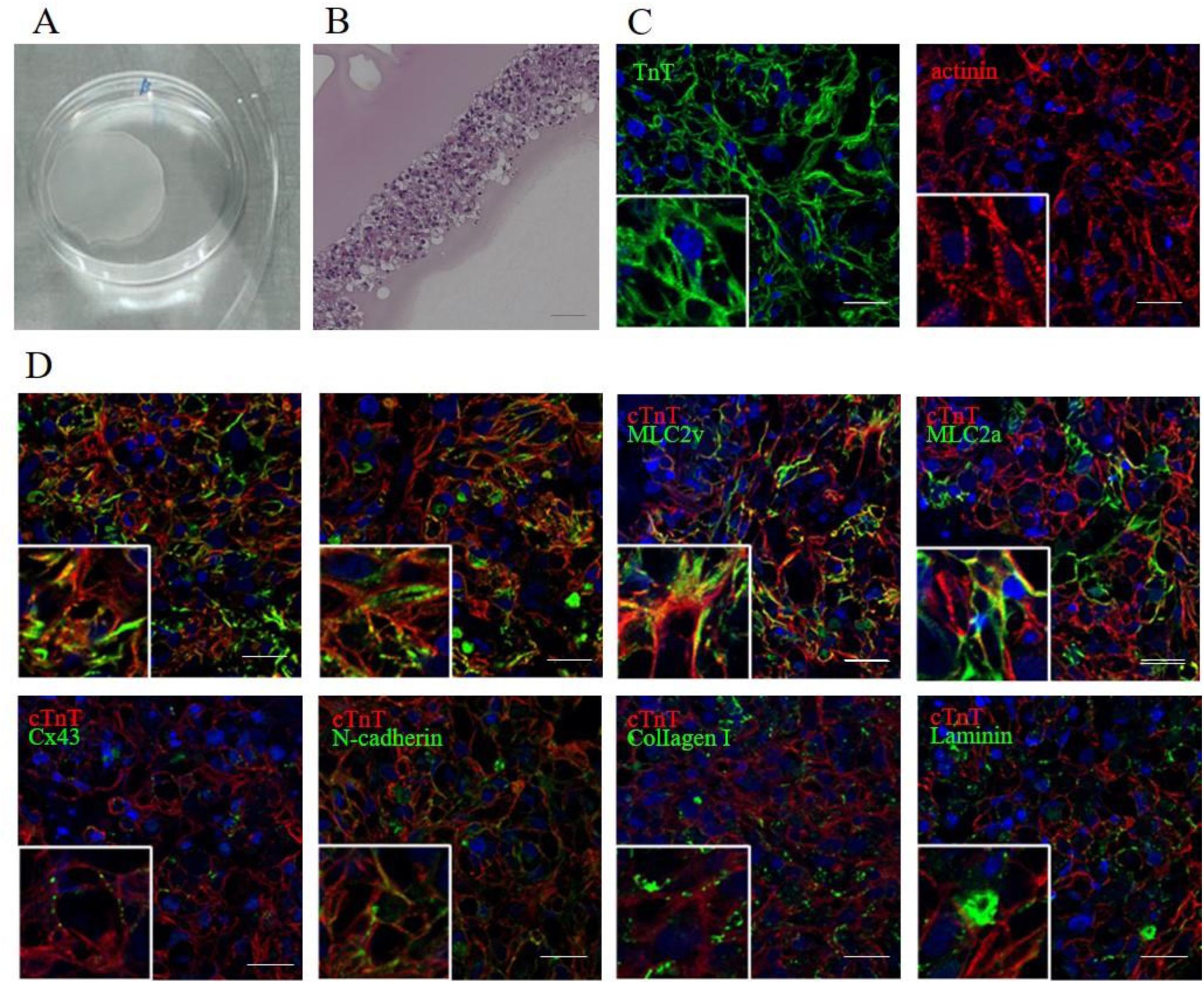
**Characteristic properties of the hiPSC-CM patch.** A: Image of a hiPSC-CM patch in a 6 cm diameter dish. B: Haematoxylin and eosin (H&E) staining of a hiPSC-CM patch. Scale bars: 50 μm (B). C, D: Representative image of an immunostained hiPSC-CM cell patch. (C) TnT (green) and actinin (red); (D) TnT (red) and αMHC, βMHC, MLC2v, MLC2a, connexin43 (cx43), N-cadherin, collagen I, and collagen III (green). Scale bars: 20 μm.

**Figure 3.**
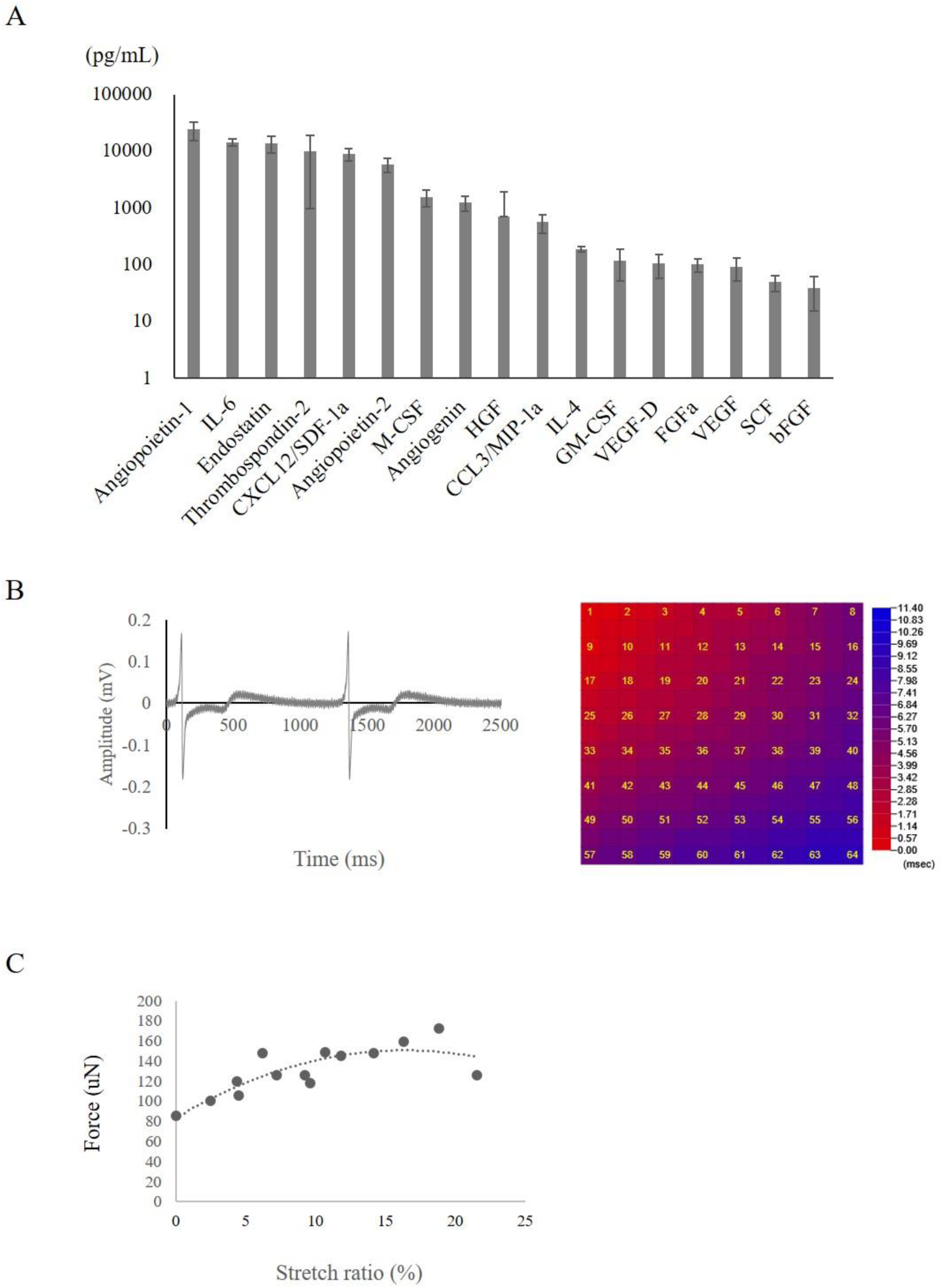
**Characterisation of the hiPSC-CM patch.** A: *In vitro* quantification of cytokines and growth factors. B: Electrophysiological properties of the hiPSC-CM patch. Extracellular field potentials were recorded using a multi-electrode array system. A representative extracellular potential waveform and a propagation map are shown. C: The contractile force of the hiPSC-CM patch was assessed using the MicroTester G2. The relationship between the contraction force and the stretch rate is shown.

### Efficacy study of iPSC-CM patches in an infarction porcine model

An *in vivo* study of the efficacy of hiPSC-CM patches was performed using a porcine infarction model (see details in Figure S7). Transthoracic echocardiography was performed before and 4, 8, and 12 weeks after hiPSC-CM patch transplantation or sham surgery. The baseline left ventricle ejection fraction (LVEF), LV end-diastolic diameter (LVDd), and LV end-systolic diameter (LVDs) did not differ significantly between the two groups (Figure 4A). LVEF was significantly greater in the hiPSC-CM patch group than in the sham group after 4 weeks (61.1 ± 5.7% vs. 46.3 ± 2.3%, *P* < 0.01), 8 weeks (60.1 ± 7.5% vs. 48.6 ± 6.1%, *P* < 0.05), and 12 weeks (63.0 ± 6.7% vs. 39.6 ± 9.8%, *P* < 0.01). LVDs was significantly smaller in the hiPSC-CM patch group than in the sham group after 4 weeks (20.7 ± 2.6 mm vs. 28.3 ± 1.3 mm, *P* < 0.01), 8 weeks (21.0 ± 3.5 mm vs. 29.0 ± 5.8 mm, *P* < 0.05), and 12 weeks (21.6 ± 4.7 mm vs. 31.0 ± 3.1 mm, *P* < 0.05), whereas LVDd did not differ significantly between the two groups.

**Figure 4.**
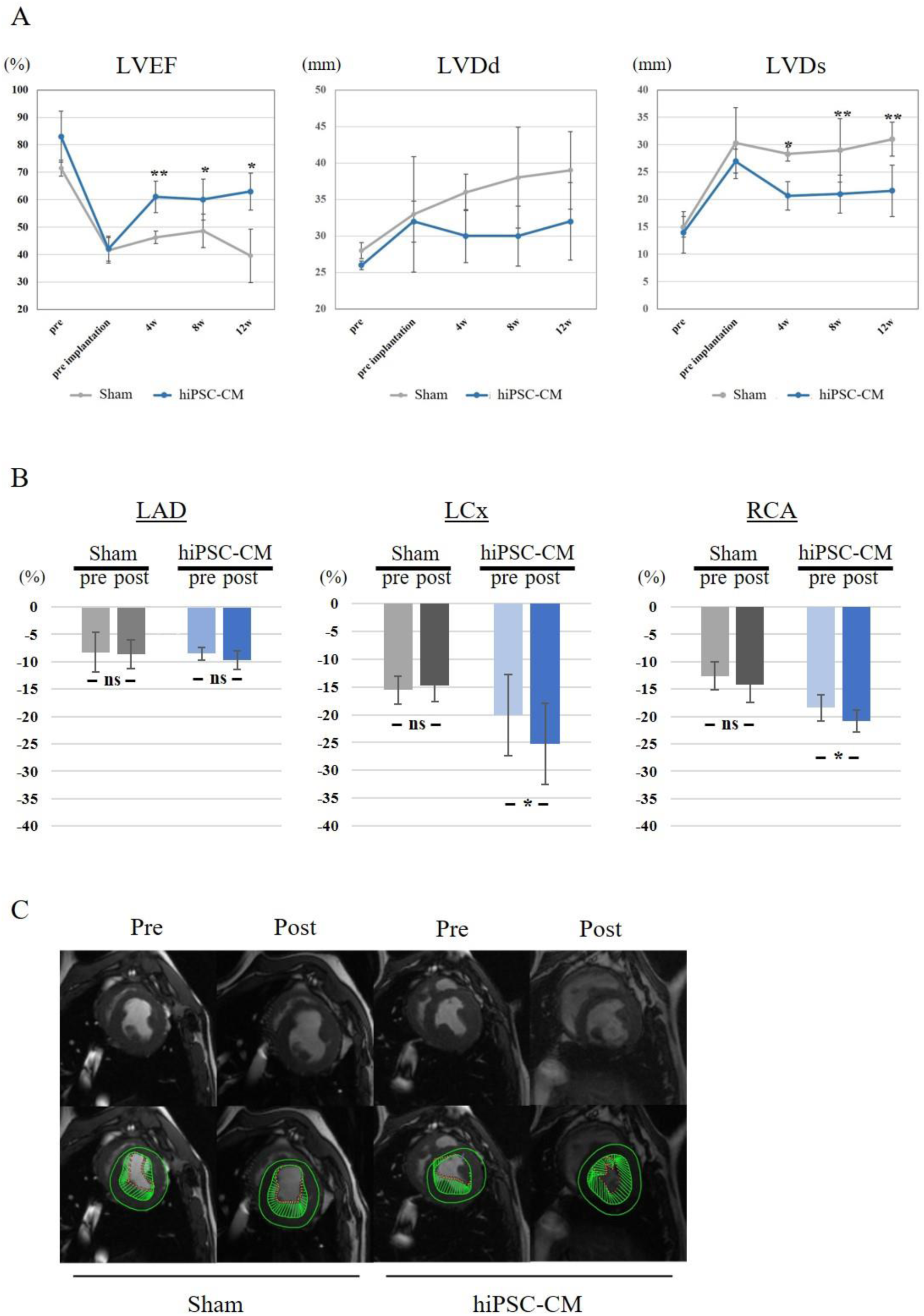
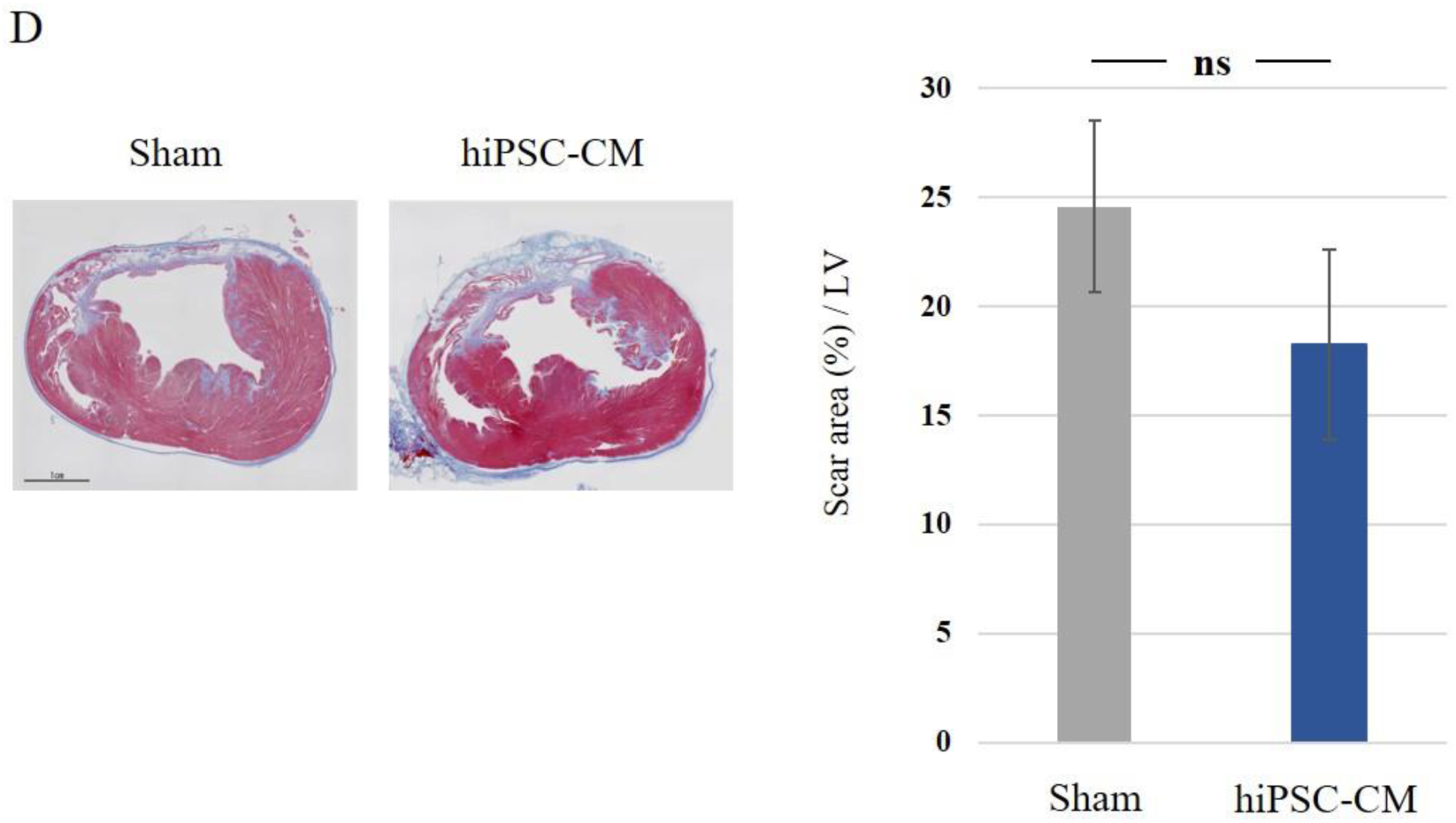
**Efficacy of hiPSC-CM patch transplantation into a porcine MI model: a pre-clinical trial.** A: Change in cardiac function after transplantation of the hiPSC-CM patch into a porcine MI model. LVEF: left ventricular ejection fraction; LVDd: left ventricular end-diastolic diameter; LVDs: left ventricular end-systolic diameter. B: Cardiac MRI was performed to compare the LV CS values at baseline and 12 weeks after treatment. C: Representative images of endocardial systolic cardiac wall motion at the papillary muscle level 12 weeks after the implantation. D: Pathological interstitial fibrosis 12 weeks after treatment. Left: Picrosirius red staining of a porcine heart. Scale bar: 1 cm. right: % area of fibrosis Data are presented as the mean ± SD. \**P* < 0.05, \*\**P* < 0.01; ns, not significant.

Cardiac MRI was performed to compare the LV circumferential strain (CS) values at pre implantation baseline and 12 weeks after implantation (Figure 4B, 4C). In the sham group, the CS of left anterior descending artery (LAD), left circumflex artery (LCx), and right coronary artery (RCA) territories did not significantly change after 12 weeks relative to the baseline values. In contrast, in the hiPSC-CM patch group, the CS of the LCx and RCA territories were significantly greater after 12 weeks than at baseline (LCx: −20.0 ± 7.3% vs. −25.5 ± 7.3%, *P* < 0.05; RCA: −18.4 ± 2.4% vs. −20.8 ± 2.1%, *P* < 0.05), whereas CS levels in the LAD territory did not significantly change.

Next, pathological interstitial fibrosis 12 weeks after treatment was assessed by Picrosirius red staining (Figure 4D). The interstitial fibrosis area did not significantly differ between the hiPSC-CM patch and sham groups (P = 0.088). An angiogram and pressure wire study was also conducted to assess treatment-induced remodelling of the coronary artery branch network (Figure 5A, 5B). The proximal LAD was completely occluded in all subjects. We defined the delta index of microvascular resistance (ΔIMR) as IMR (post-implantation) - IMR (pre-implantation). ΔIMR in the LCx territory (infarcted-border region), such as the posterolateral branch (PL) (-20.0 ± 28.2 versus 38.4 ± 12.2, *P* < 0.05) and obtuse marginal branch (OM) (-17.0 ± 11.1 versus 34.3 ± 23.7, *P* < 0.05) branches, was significantly lower in the hiPSC-CM patch group than in the sham group. The IMR in the RCA territory (infarcted-remote region) was not significantly different between the two groups.

**Figure 5.**
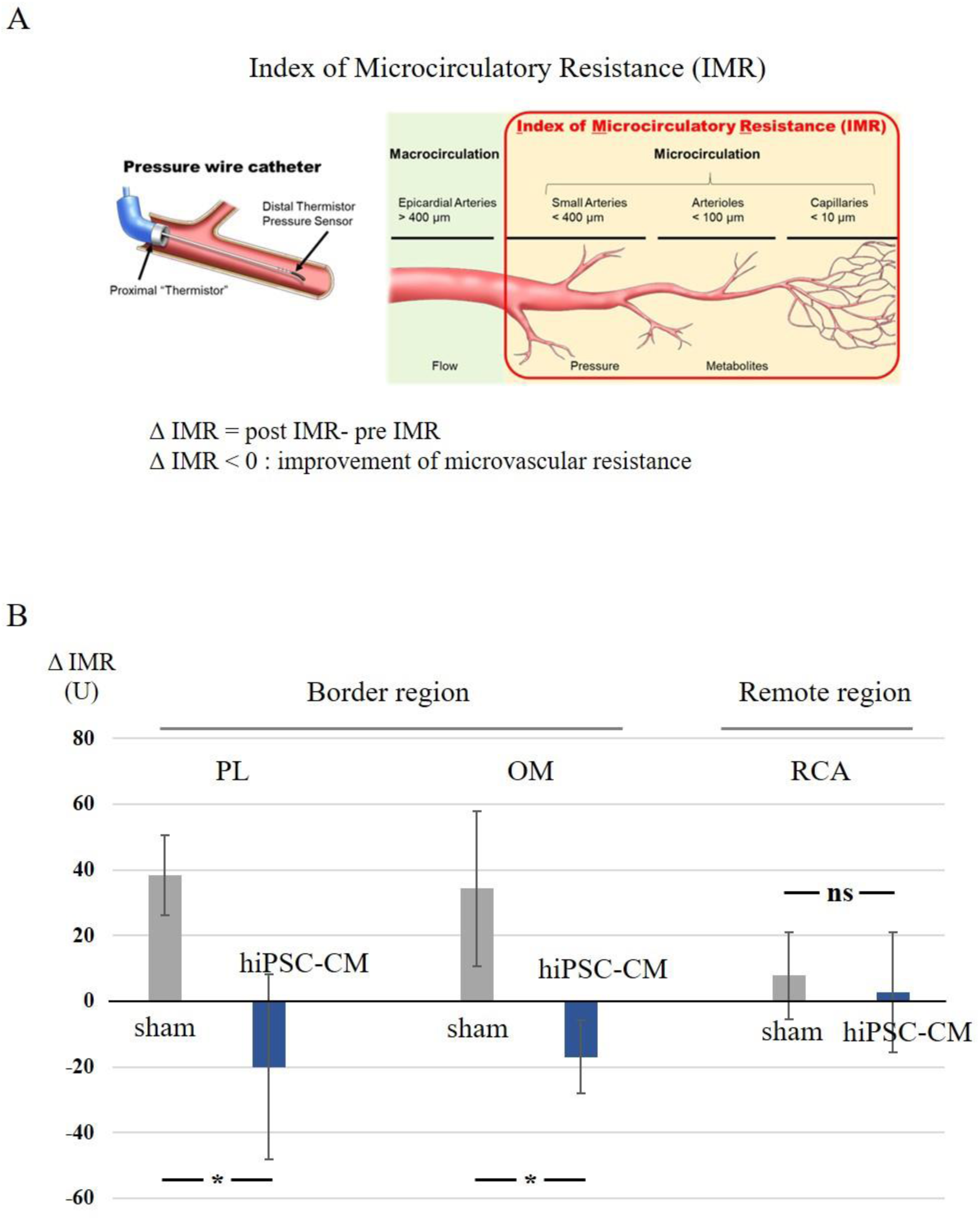
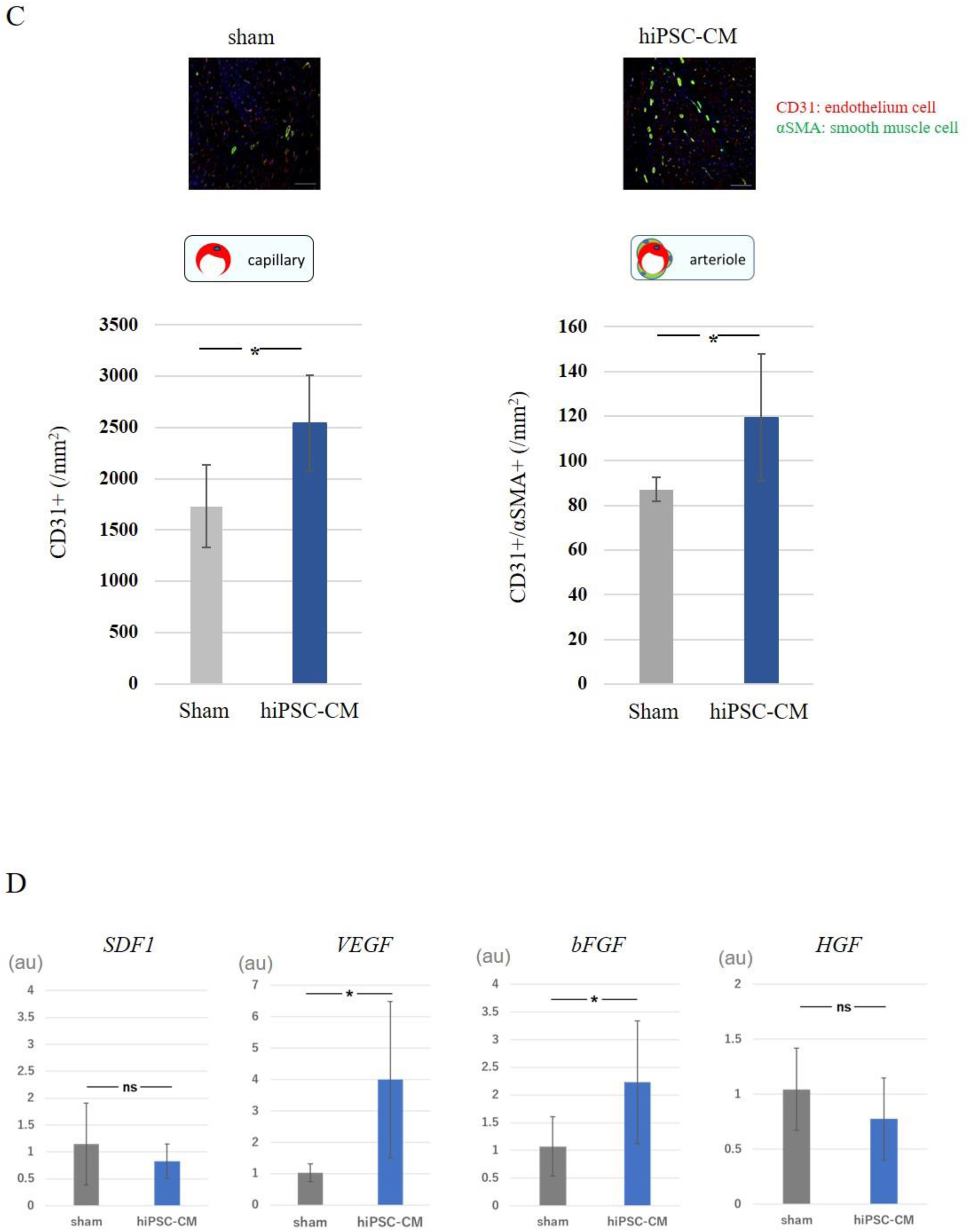
**Change in IMR and cytokine expression after transplantation of the hiPSC-CM patch into a porcine MI model.** A, B: Schematic representation of IMR. ΔIMP was defined as IMR (post-implantation) - IMR (pre-implantation). C: upper panel, representative image of capillaries and arterioles immunostained with CD31 (red) and αSMA (green); lower panel, quantification of the number of CD31- and αSMA-positive cells. Scale bar: 100 μm. D: Gene expression of proangiogenic factors in the infarct-border region 4 weeks after treatment. Data are presented as the mean ± SD. \**P* < 0.05; ns, not significant.

Vascular density 12 weeks after treatment was assessed by immunohistochemistry for CD31 and αSMA (Figure 5C). The density of CD31-positive capillaries and CD31/αSMA double-positive arterioles was significantly greater in the hiPSC-CM patch group than in the sham group (2542.2 ± 465.3/mm^2^ vs. 1732.4 ± 405.2/mm^2^, *P* < 0.05; 119.4 ± 28.5/mm^2^ vs. 87.2 ± 5.5/mm^2^, *P* < 0.05, respectively; Figure 5C). Moreover, gene expression of proangiogenic cytokines in the infarct-border region 4 weeks after treatment was also assessed. VEGF and basic fibroblast growth factor (bFGF) expression levels were significantly higher in the infarct-border region of the hiPSC-CM patch group than in the sham group (VEGF: 4.00 ± 2.48 vs. 1.03 ± 0.28, *P* < 0.05; bFGF: 2.23 ± 1.10 vs. 1.07 ± 0.53), whereas SDF-1 and HGF expression did not significantly differ between the two groups (Figure 5D).

To assess the effects of the hiPSC-CM patch on the electrophysiology of the myocardium, the patch and sham groups were monitored via 24-hr Holter electrocardiography 7 days before and 0, 1, 2, 3, 7, 14, 28, 42, 56, 70, and 84 days after transplantation (Table S6). No lethal arrhythmias such as ventricular tachycardia and ventricular fibrillation were observed during the study period. In addition, tumour formation was not detected in the three months after implantation for both groups.

### Safety assessment for tumorigenicity and toxicity

To evaluate the safety of hiPSC-CM patches, general toxicity and tumourigenicity tests were performed. To assess general toxicity, a single hiPSC-CM patch was applied to the heart surface of male and female NOG mice and observed for 4 weeks. The general condition, body weight, results of haematology and blood biochemistry tests, and pathological assessment were recorded. No significant toxic changes were observed following application of the hiPSC-CM patch (Table S7).

To assess tumourigenicity, we monitored the animals for teratomas and malignant tumours caused by residual undifferentiated hiPSCs and malignant transformed cells, respectively, using *in vitro* and *in vivo* assays previously reported ^15^ for the detection of potential tumorigenic cells in hiPSC-CMs. *In vitro*, we performed assays for cell growth and soft agar colony formation—a highly sensitive method for detecting malignant transformed cells—to identify cells that have undergone malignant transformation during hiPSC culture and cardiomyocyte differentiation. The growth rate of hiPSC-CMs during P5 was significantly lower than during initial passage (−0.12 ± 0.04 vs. 0.14 ± 0.03 doubling/day, *P* < 0.01), indicating that hiPSC-CMs contain no abnormal, overgrowing cells (Figure 6A). The soft agar colony formation assay showed no growth of hiPSC-CMs (Figure 6B). To detect teratoma-forming cells in the hiPSC-CM patch, the expression of Lin28A—a marker of undifferentiated hiPSCs—, was examined *in vitro* (Figure S2C). Lin28A expression was below the limit of determination in hiPSC-CMs 30 days after cardiac differentiation. Finally, we assessed tumourigenicity *in vivo* and found that no teratomas or malignant tumours formed in the heart for 16 weeks after transplantation of hiPSC-CM patches (Figure 6C, Table S7).

**Figure 6.**
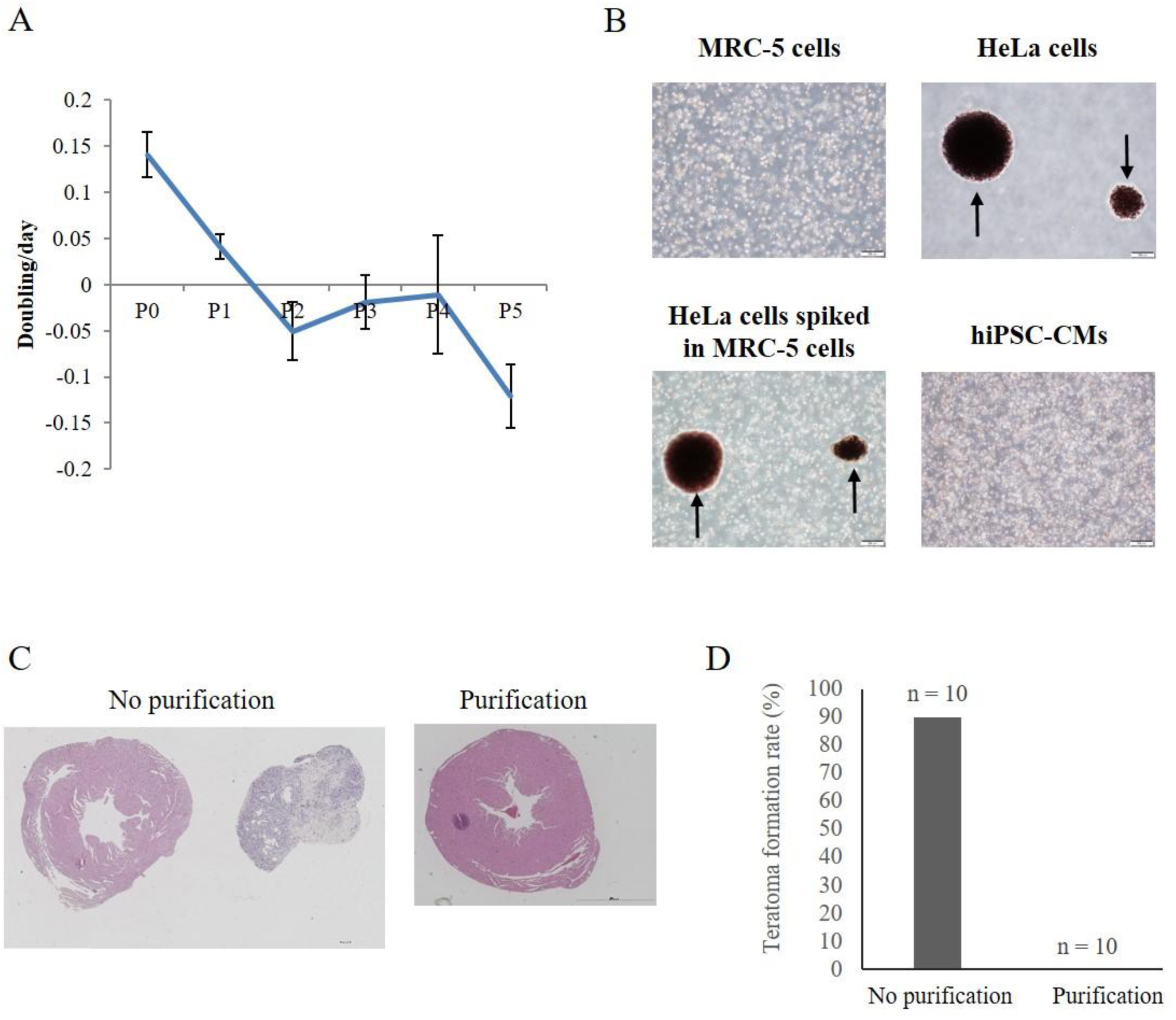
**Detection of tumourigenic cells *in vitro* and *in vivo*.** A: Cell growth assay of each passage. B: Soft agar colony formation assay. Phase contrast micrographs of MRC-5 cells, HeLa cells, HeLa cells spiked into MRC-5 cells, and hiPSCs-CMs cultured on soft agar medium for 21 days. Arrows indicate colonies. Scale bar: 200 μm. C, D: Tumourigenicity was evaluated through transplantation of non-purified or purified hiPSC-CM patches into the left ventricular surface of immunodeficient NOG mice. (C) Representative H&E staining of teratoma. Scale bar: 1000 µm (left panel) and 2000 µm (right panel). (D) Quantification of the rate of teratoma formation (no purification group: hiPSC-CM patches without purification and elimination of residual undifferentiated hiPSCs, n = 10; purification group: hiPSC-CM patches with purification and elimination of residual undifferentiated hiPSCs, n = 10).

In addition to residual undifferentiated cells or malignant transformed cells, critical genomic changes and the survival of foreign genes in hiPSCs may lead to tumour formation in hiPSC-CMs. We thus performed WGS and WES of hiPSCs, hiPSC-CMs, and hiPSC-CM patches. No single nucleotide variants (SNV)/indels were found for the cancer-related genes listed in the Catalogue of Somatic Mutations in Cancer (COSMIC) ^16^, the Cancer Gene Census (v79) ^17^, and Shibata’s list^18^. In addition, called mutations were not registered in the Human Gene Mutation Database (HGMD) Pro database (2016.4) ^19^. We calculated variant allele frequencies (VAFs; Table 1) at SNV/indel positions found by Genomon^20^ and Genomon2^21^ in the context of WGS and WES data. We also investigated CNVs based on WGS and SNP array data. No CNVs were found on exons. The genome analyses are summarised in Table 2.

**Table 1.** Whole genome sequencing results for hiPSCs (MCB), hiPSC-CMs, and hiPSC-CM patches.

| Sample | # of SNVs/indels on CDS/splicing sites |  | # of SNVs/indels on CDS/splicing sites (Census/Shibata's list) | # of CNVs on exons | # of CNVs on exons (Census/Shibata's list) |
| --- | --- | --- | --- | --- | --- |
|  | WGS | WES | WGS/WES | WGS/SNP array | WGS/SNP array |
| MCB | 15 | 11 | 0 | 0 | 0 |
| MCB (expansion culture) | 14 | 11 | 0 | 0 | 0 |
| hiPSC-CMs | 14 | 12 | 0 | 0 | 0 |
| hiPSC-CM patch | 14 | 11 | 0 | 0 | 0 |

hiPSC-CM, human induced pluripotent stem cell-derived cardiomyocyte; SNV, single nucleotide variant; Indels,
insertions/deletions; CNV, copy number variation; MCB, master cell bank; WGS, whole genome sequencing;
WES, whole exome sequencing; SNP, single-nucleotide polymorphism.

**Table 2.**
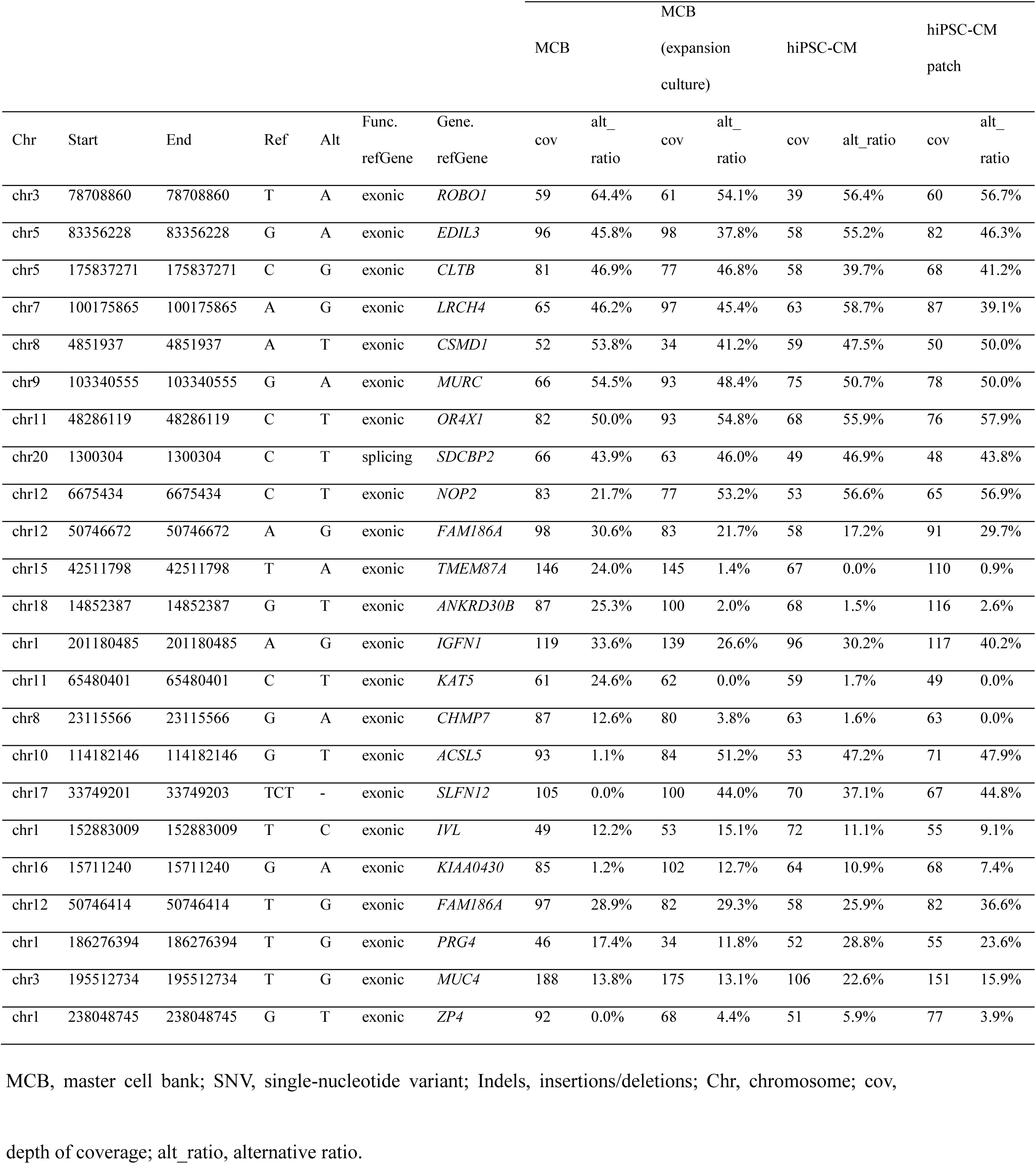
Variant allele frequencies as per the called SNVs/indels.

## Discussion

In this paper, we reported the biological characterisation, efficacy, and safety of clinical grade hiPSC-CM patches as part of a pre-clinical study. In hiPSC-CMs, the cTnT-positive rate ranged from 60 to 80%, while Lin28A, which represents undifferentiated hiPSCs, was below the limit of quantitation. The safety of hiPSC-CM patches for clinical applications was confirmed through genomic analysis; *in vitro* cell growth, soft agar colony formation, and undifferentiated cell assays; *in vivo* tumourigenicity and general toxicity tests using immunodeficient mice; and an arrhythmia test using a porcine model. Finally, an efficacy study using a porcine myocardial infarction (MI) model demonstrated that hiPSC-CM patches ameliorated the distressed myocardium in terms of improved cardiac function and angiogenesis.

Although the teratogenicity of undifferentiated hiPSCs can be adequately determined through Lin28A expression and controlled by brentuximab vedotin, which induces apoptosis of CD30-positive undifferentiated hiPSCs, the tumourigenicity of malignant transformed cells in hiPSC-CMs is still a major obstacle to clinical translation^15,, 22^. Thus, we performed a genomic analysis to confirm the tumourigenicity of hiPSC-CM patches, and although no abnormal mutations were found, it remains undetermined which genomic mutations should be checked to ensure safety other than those reported to be tumourigenic in oncogenomic databases such as COSMIC^16^ and Shibata’s list^18^. Various genetic mutations have been identified in living cells, and it is difficult to determine whether all mutations of potentially tumourigenic genes result in oncogenesis or whether some well-known tumourigenicity-associated genes such as c-myc can be ignored. Consequently, the relationship between genomic abnormalities and tumourigenicity in cell therapy has not been fully elucidated, and further studies are thus warranted to confirm the safety of these cells in terms of tumourigenicity. Nevertheless, in immunodeficient NOG mice, no malignant transformed cells or tumourigenicity were observed.

We need to reveal both the tumourigenicity of residual undifferentiated hiPSCs and verify the presence of tumourigenic cells by transformation of hiPSC-CM constructs. Lin28A expression levels have been reported to correlate with the frequency of tumour formation in NOG mice ^15^. Quantifying Lin28A levels may thus be an alternative to tumourigenicity tests of NOG mice when validating the safety of hiPSC-CMs in clinical applications.

In this study, no lethal arrhythmia was observed when hiPSC-CM patches were transplanted on the heart surface of pigs. In contrast, Shiba et al.^5^ reported that arrhythmias are markedly increased in the early phase after hiPSC-CM transplantation with a needle. Liu et al.^23^ created artificial myocardial tissue using hiPSC-CMs, and when the tissue was injured with a needle, they observed a rotating electrical pulse reminiscent of lethal arrhythmia around the injury site. This abnormal conduction was restored only after applying a cardiomyocyte patch on the injured site. Menache et al.^24^ also detected the occurrence of lethal arrhythmia when myoblasts were injected into myocardial tissue with a needle. This discrepancy between our results and previous studies may have resulted from the method by which cells were introduced, such as needle injection. Thus, the hiPSC-CM patch avoids tumourigenicity and arrhythmogenicity, indicating it may be a safe method of delivering cells. A clinical study is necessary to guarantee the safety of this technique.

An important propertyof hiPSC-CM patches is whether the transplanted cardiac patches can be electrically integrated with a recipient heart with a low number of cardiomyocytes. Previous studies have shown that the transplanted cardiomyocyte patches contract and relax in synchrony with the recipient’s heart^4, 25^. It was also demonstrated that cardiomyocyte patches exhibit excellent cardiogenic properties, cardiomyogenesis potential in the failed heart, and angiogenic ability^26–29^. Further studies are needed to elucidate how many cardiomyocytes can directly provide contractile force to the diseased heart and the fundamental mode of action of this treatment.

Functional recovery of the heart may also depend on angiogenesis, which may have a positive impact on hibernating myocardium in the recipient myocardium^26–29^. Histological analysis suggests that it is characterised by the recognition of functional blood vessels with smooth muscle cells lining vascular endothelial cells^26–29^. In particular, cytokines of the angiopoietin family may be enriched *in vitro*. Angiopoietins have been reported to greatly contribute to the maturation of blood vessels. In general, ischaemic cardiomyopathy is characterised by myocardial ischaemia arising from the disruption and stenosis of the vasculature network in coronary arteries^26–29^. Here, the heart failure animal model transplanted with hiPSC-CM patches demonstrated low peripheral coronary vascular resistance, which was likely due to the maturation of blood vessels and opening of the occluded peripheral vascular network.

In conclusion, we conducted a proof-of-concept, pre-clinical study of hiPSC-CM patches, which demonstrated promising feasibility, safety, and efficacy. We recommend further translational research by conducting a clinical trial of allogenic hiPSC-CM patches for patients with ischaemic heart failure.

## Acknowledgements

We appreciate the assistance of Professor Shinya Yamanaka at the Center for iPS Cell Research and Application, Kyoto University, who kindly provided the hiPSCs. The authors wish to thank Professor Tetsuya Shimizu and Associate Professor Katsuhisa Matsuura at Tokyo Women’s Medical University and Associate Professor Yoshinori Yoshida at Kyoto University for their collaboration during the early stages of this work. We also thank Fumiya Ohashi, Kenji Oyama, and Isamu Matsuda at Terumo. Co. and Kousuke Torigata at Daiichi Sankyo. Co. for their technical assistance with the experiments.

## Data availability statement

The data underlying this article cannot be shared publicly due to the privacy of individuals that participated in the study. The data will be shared on reasonable request to the corresponding author.

## References

1. Yoshida Y, Yamanaka S. Induced Pluripotent Stem Cells 10 Years Later: For Cardiac Applications. Circ Res 2017;120(12):1958–1968.

2. Mandai M, Watanabe A, Kurimoto Y, Hirami Y, Morinaga C, Daimon T, Fujihara M, Akimaru H, Sakai N, Shibata Y, Terada M, Nomiya Y, Tanishima S, Nakamura M, Kamao H, Sugita S, Onishi A, Ito T, Fujita K, Kawamata S, Go MJ, Shinohara C, Hata KI, Sawada M, Yamamoto M, Ohta S, Ohara Y, Yoshida K, Kuwahara J, Kitano Y, Amano N, Umekage M, Kitaoka F, Tanaka A, Okada C, Takasu N, Ogawa S, Yamanaka S, Takahashi M. Autologous Induced Stem-Cell-Derived Retinal Cells for Macular Degeneration. N Engl J Med 2017;376(11):1038–1046.

3. Doi D, Magotani H, Kikuchi T, Ikeda M, Hiramatsu S, Yoshida K, Amano N, Nomura M, Umekage M, Morizane A, Takahashi J. Pre-clinical Study of Induced Pluripotent Stem Cell-Derived Dopaminergic Progenitor Cells for Parkinson’s Disease. Nat Commun 2020;11(1):3369.

4. Higuchi T, Miyagawa S, Pearson JT, Fukushima S, Saito A, Tsuchimochi H, Sonobe T, Fujii Y, Yagi N, Astolfo A, Shirai M, Sawa Y. Functional and Electrical Integration of Induced Pluripotent Stem Cell-Derived Cardiomyocytes in a Myocardial Infarction Rat Heart. Cell Transplant 2015;24(12):2479–2489.

5. Shiba Y, Gomibuchi T, Seto T, Wada Y, Ichimura H, Tanaka Y, Ogasawara T, Okada K, Shiba N, Sakamoto K, Ido D, Shiina T, Ohkura M, Nakai J, Uno N, Kazuki Y, Oshimura M, Minami I, Ikeda U. Allogeneic Transplantation of Ips Cell-Derived Cardiomyocytes Regenerates Primate Hearts. Nature 2016;538(7625):388–391.

6. Tabei R, Kawaguchi S, Kanazawa H, Tohyama S, Hirano A, Handa N, Hishikawa S, Teratani T, Kunita S, Fukuda J, Mugishima Y, Suzuki T, Nakajima K, Seki T, Kishino Y, Okada M, Yamazaki M, Okamoto K, Shimizu H, Kobayashi E, Tabata Y, Fujita J, Fukuda K. Development of a Transplant Injection Device for Optimal Distribution and Retention of Human Induced Pluripotent Stem Cell-Derived Cardiomyocytes. J Heart Lung Transplant 2019;38(2):203–214.

7. Memon IA, Sawa Y, Fukushima N, Matsumiya G, Miyagawa S, Taketani S, Sakakida SK, Kondoh H, Aleshin AN, Shimizu T, Okano T, Matsuda H. Repair of Impaired Myocardium by Means of Implantation of Engineered Autologous Myoblast Sheets. J Thorac Cardiovasc Surg 2005;130(5):1333–1341.

8. Cunningham JJ, Ulbright TM, Pera MF, Looijenga LH. Lessons from Human Teratomas to Guide Development of Safe Stem Cell Therapies. Nat Biotechnol 2012;30(9):849–857.

9. Yasuda S, Kusakawa S, Kuroda T, Miura T, Tano K, Takada N, Matsuyama S, Matsuyama A, Nasu M, Umezawa A, Hayakawa T, Tsutsumi H, Sato Y. Tumorigenicity-associated Characteristics of Human iPS Cell Lines. PLoS One 2018;13(10):e0205022.

10. Yamanaka S. Pluripotent Stem Cell-based Cell Therapy––Promise and Challenges. Cell Stem Cell 2020;27(4):523–531.

11. Okita K, Matsumura Y, Sato Y, Okada A, Morizane A, Okamoto S, Hong H, Nakagawa M, Tanabe K, Tezuka K, Shibata T, Kunisada T, Takahashi M, Takahashi J, Saji H, Yamanaka S. A More Efficient Method to Generate Integration-Free Human Ips Cells. Nat Methods 2011;8(5):409–412.

12. Wang K, Li M, Hadley D, Liu R, Glessner J, Grant SF, Hakonarson H, Bucan M. PennCNV: An Integrated Hidden Markov Model Designed for High-resolution Copy Number Variation Detection in Whole-genome SNP Genotyping Data. Genome Res 2007;17(11):1665–1674.

13. González JR, Rodríguez-Santiago B, Cáceres A, Pique-Regi R, Rothman N, Chanock SJ, Armengol L, Pérez-Jurado LA. A Fast and Accurate Method to Detect Allelic Genomic Imbalances Underlying Mosaic Rearrangements Using SNP Array Data. BMC Bioinformatics 2011;12:166.

14. Gogarten SM, Bhangale T, Conomos MP, Laurie CA, McHugh CP, Painter I, Zheng X, Crosslin DR, Levine D, Lumley T, Nelson SC, Rice K, Shen J, Swarnkar R, Weir BS, Laurie CC. GWASTools: An R/Bioconductor Package for Quality Control and Analysis of Genome-Wide Association Studies. Bioinformatics 2012;28(24):3329–3331.

15. Ito E, Miyagawa S, Takeda M, Kawamura A, Harada A, Iseoka H, Yajima S, Sougawa N, Mochizuki-Oda N, Yasuda S, Sato Y, Sawa Y. Tumorigenicity Assay Essential for Facilitating Safety Studies of hiPSC-derived Cardiomyocytes for Clinical Application. Sci Rep 2019;9(1)1881.

16. Tate JG, Bamford S, Jubb HC, Sondka Z, Beare DM, Bindal N, Boutselakis H, Cole CG, Creatore C, Dawson E, Fish P, Harsha B, Hathaway C, Jupe SC, Kok CY, Noble K, Ponting L, Ramshaw CC, Rye CE, Speedy HE, Stefancsik R, Thompson SL, Wang S, Ward S, Campbell PJ, Forbes SA. COSMIC: The Catalogue Of Somatic Mutations in Cancer. Nucleic Acids Res 2019;47(D1):D941–D947.

17. Sondka Z, Bamford S, Cole CG, Ward SA, Dunham I, Forbes SA. The COSMIC Cancer Gene Census: Describing Genetic Dysfunction Across all Human Cancers. Nat Rev Cancer 2018;18(11):696–705.

18. Nakahata T, Okano H. Current Perspective on Evaluation of Tumorigenicity of Cellular and Tissue-based Products Derived from Induced Pluripotent Stem Cells (iPSCs) and iPSCs as Their Starting Materials (provisional translation). Pharmaceuticals and Medical Devices Agency. http://www.pmda.go.jp/files/000152599.pdf

19. Stenson PD, Mort M, Ball EV, Evans K, Hayden M, Heywood S, Hussain M, Phillips AD, Cooper DN. The Human Gene Mutation Database: Towards a Comprehensive Repository of Inherited Mutation Data For Medical Research, Genetic Diagnosis and Next-generation Sequencing Studies. Hum Genet 2017;136(6):665–677.

20. Yoshida K, Sanada M, Shiraishi Y, Nowak D, Nagata Y, Yamamoto R, Sato Y, Sato-Otsubo A, Kon A, Nagasaki M, Chalkidis G. Frequent Pathway Mutations of Splicing Machinery in Myelodysplasia. Nature 2011;478(7367):64–69.

21. Shiraishi Y, Sato Y, Chiba K, Okuno Y, Nagata Y, Yoshida K, Shiba N, Hayashi Y, Kume H, Homma Y, Sanada M, Ogawa S, Miyano S. An Empirical Bayesian Framework for Somatic Mutation Detection from Cancer Genome Sequencing Data. Nucleic Acids Res 2013;41(7):e89.

22. Sougawa N, Miyagawa S, Fukushima S, Kawamura A, Yokoyama J, Ito E, Harada A, Okimoto K, Mochizuki-Oda N, Saito A, Sawa Y. Immunologic Targeting of CD30 Eliminates Tumourigenic Human Pluripotent Stem Cells, Allowing Safer Clinical Application of Hipsc-Based Cell Therapy. Sci Rep 2018;8(1):3726.

23. Li J, Minami I, Shiozaki M, Yu L, Yajima S, Miyagawa S, Shiba Y, Morone N, Fukushima S, Yoshioka M, Li S, Qiao J, Li X, Wang L, Kotera H, Nakatsuji N, Sawa Y, Chen Y, Liu L. Human Pluripotent Stem Cell-Derived Cardiac Tissue-like Constructs for Repairing the Infarcted Myocardium. Stem Cell Reports 2017;9(5):1546–1559.

24. Menasché P, Alfieri O, Janssens S, McKenna W, Reichenspurner H, Trinquart L, Vilquin JT, Marolleau JP, Seymour B, Larghero J, Lake S, Chatellier G, Solomon S, Desnos M, Hagège AA. The Myoblast Autologous Grafting in Ischemic Cardiomyopathy (MAGIC) Trial: First Randomized Placebo-controlled Study Of Myoblast Transplantation. Circulation 2008;117(9):1189–1200.

25. Iseoka H, Miyagawa S, Fukushima S, Saito A, Masuda S, Yajima S, Ito E, Sougawa N, Takeda M, Harada A, Lee JK, Sawa Y. Pivotal Role of Non-cardiomyocytes in Electromechanical and Therapeutic Potential of Induced Pluripotent Stem Cell-Derived Engineered Cardiac Tissue. Tissue Eng Part A 2018;24(3-4):287–300.

26. Kawamura M, Miyagawa S, Miki K, Saito A, Fukushima S, Higuchi T, Kawamura T, Kuratani T, Daimon T, Shimizu T, Okano T, Sawa Y. Feasibility, Safety, and Therapeutic Efficacy of Human Induced Pluripotent Stem Cell-Derived Cardiomyocyte Sheets in a Porcine Ischemic Cardiomyopathy Model. Circulation 2012;126(11:Suppl 1):S29–S39.

27. Kawamura M, Miyagawa S, Fukushima S, Saito A, Miki K, Ito E, Sougawa N, Kawamura T, Daimon T, Shimizu T, Okano T, Toda K, Sawa Y. Enhanced Survival of Transplanted Human Induced Pluripotent Stem Cell-derived Cardiomyocytes by the Combination of Cell Sheets with the Pedicled Omental Flap Technique in a Porcine Heart. Circulation 2013;128(11 Suppl 1):S87–S94.

28. Kawamura M, Miyagawa S, Fukushima S, Saito A, Miki K, Funakoshi S, Yoshida Y, Yamanaka S, Shimizu T, Okano T, Daimon T, Toda K, Sawa Y. Enhanced Therapeutic Effects of Human iPS Cell Derived-Cardiomyocyte by Combined Cell-Sheets with Omental Flap Technique in Porcine Ischemic Cardiomyopathy Model. Sci Rep 2017;7(1):8824.

29. Ishida M, Miyagawa S, Saito A, Fukushima S, Harada A, Ito E, Ohashi F, Watabe T, Hatazawa J, Matsuura K, Sawa Y. Transplantation of Human-induced Pluripotent Stem Cell-derived Cardiomyocytes is Superior to Somatic Stem Cell Therapy for Restoring Cardiac Function and Oxygen Consumption in a Porcine Model of Myocardial Infarction. Transplantation 2019;103(2):291–298.

