## Supplementary_data for "Evaluation of the Efficacy and Safety of a Clinical Grade Human Induced Pluripotent Stem Cell-Derived Cardiomyocyte Patch: A Pre-Clinical Study"

<Translational Research article>

### Supplementary files

#### Supplementary methods

##### Clinical grade hiPSCs

The clinical grade hiPS cell line (QHJI14s04) was established using peripheral blood mononuclear cells collected from a healthy HLA homozygous (HLA-A, HLA-B, HLA-C, HLA-DRB1, HLA-DPB1, HLA-DQB1) donor, showing the most frequent haplotype in the Japanese population. QHJI14s04 was generated using episomal plasmids[1] (pCE-hSK, pCE-hUL, pCE-hOCT3/4, pCE-mp53DD, pCXB-EBNA1; Table S2) and maintained using a feeder-free and xeno-free culture system[2] in the cell processing centre of the Center for iPS Cell Research and Application (Kyoto University, Kyoto, Japan). We performed tests to ensure sterility and identity, as well as characterisation tests including the evaluation of cell morphology, viability, vector retention, expression of undifferentiated markers, and genomic analysis (Table S3).

##### Cardiomyogenic differentiation, purification, and elimination of residual undifferentiated hiPSCs

Cardiomyogenic differentiation of QHJI14s04 cells was induced as previously described[3]. We generated embryoid bodies (EBs) from hiPSCs, and cultured them in 100 mL bioreactors (Able corp., Tokyo, Japan) with various recombinant proteins and chemical compounds for cardiomyogenic differentiation. After differentiation, cardiomyocyte aggregates were cultured in glucose-free Dulbecco's modified Eagle's medium (DMEM; Nacalai Tesque, Kyoto, Japan) for purification. Afterward, cardiomyocytes were dissociated as previously described[4]. The dissociated cells were cultured in DMEM supplemented with 10% foetal bovine serum (FBS; Sigma-Aldrich, St. Louis, MO) and brentuximab vedotin (ADCETRIS™, Takeda, Osaka, Japan) for the elimination of residual undifferentiated cells[5]. The cells were suspended in a cell banker (Nippon Genetics, Tokyo, Japan) and frozen using a programme freezer (FZ2000; STREX Inc. Osaka, Japan).

**Flow cytometry**

After fixation in Fixation/Permeabilization solution (BD Biosciences, Franklin Lakes, NJ), cells were labelled with the following antibodies: anti-cTNT (1:300; sc-20025; Santa Cruz Biotechnology, Dallas, TX), anti-αSMA (1:100; ab32575; Abcam, Cambridge, UK), anti-vimentin (1:100; ab92547; Abcam), and anti-CD31 (1:10; 561654; BD Biosciences), followed by incubation with fluorescently conjugated secondary antibodies. Cell populations were resolved using the FACSCanto II system (BD Biosciences). Data were analysed using Diva (BD Biosciences) and FlowJo (TreeStar Inc., Ashland, OR) software.

**RNA isolation and quantitative polymerase chain reaction (qPCR)**

Total RNA was extracted from the cells and heart tissues using the RNeasy Mini Kit (Qiagen, Hilden, Germany) and reverse-transcribed into cDNA using the Super Script VILO cDNA Synthesis kit (Thermo Fisher Scientific, Waltham, MA). qPCR was performed with the ViiA 7™ Real-Time PCR or ABI PRISM 7700 systems (Applied Biosystems, Foster City, CA) using either SYBR Green (Applied Biosystems) or TaqMan™ probes (Applied Biosystems). The primer sequences used in this study are listed in Table S8. Each sample was analysed in triplicate. Expression of target genes was normalised to that of the glyceraldehyde-3-phosphate dehydrogenase (*GAPDH*) as control. Relative gene expression was determined using the delta-delta CT method. The data were analysed using the software provided with the ViiA 7™ Real-Time PCR or ABI PRISM 7700 systems.

**PCR arrays**

Total RNA was extracted from the hiPSCs and hiPSC-CMs using the RNeasy Mini Kit. Human adult and foetal heart total RNA was purchased from Takara Bio (Kusatsu, Japan). For the analysis of human stem cell-associated genes, cDNA was synthesised using the $RT^2$ First Strand Kit (Qiagen). The ViiA 7™ Real-Time PCR system was used to run the $RT^2$ Profiler™ PCR Array Human Stem Cell array (Qiagen). Cluster analysis was performed using the online $RT^2$ Profiler™ PCR Array software provided by SABiosciences (freely available from https://dataanalysis.qiagen.com/pcr/arrayanalysis.php?target=plothome). For the analysis of cardiac differentiation-associated genes, cDNA was synthesised using the Super Script VILO cDNA Synthesis Kit (Thermo Fisher Scientific) and gene expression was analysed using the TaqMan™ Array Human Cardiomyocyte Differentiation by BMP Receptors (Thermo Fisher Scientific).

**Single-cell preparation and single-cell RNA-seq**

The single-cell sequencing library was generated using the ICELL8 cx platform (Takara Bio). In brief, isolated cells were stained with a mixture of Hoechst 33342 and propidium iodide (R37610; Thermo Fischer Scientific) according to manufacturer's instructions. After staining, cells were washed with PBS and counted with a haemocytometer. The cell suspension was then pipetted into a 384-well plate, which was then dispensed on the ICELL8 cx system into 250 nL chips (640199; Takara Bio). Imaging and analysis of the nano-wells were carried out using the CellSelect Software, and single live cells, defined by Hoechst-positive and propidium iodide-negative staining, were selected. After dispensing the RT-PCR reaction MasterMix (640167; Takara Bio) into the selected nano-wells, the chip was sealed, centrifuged, and placed into a Chip Cycler (Bio-Rad Laboratories, Hercules, CA) for reverse transcription and full-length cDNA synthesis, following the manufacturers' protocols. The resulting cDNAs were purified, and fragments up to 300–350 bp were removed using 0.6X Agencourt AMPure XP beads (A63880; Beckman Coulter, Brea, CA). One nanogram of pooled cDNA was used as input to generate a sequencing library using the Nextera XT DNA sample preparation Kit (FC-131-1024; Illumina, San Diego, CA), following the manufacturers' protocols. Libraries were sequenced on the HiSeq 3000 sequencer (Illumina) using the 100 bp paired-end sequencing protocol.

For single-cell RNA-seq analysis, raw reads were processed using the mappa/hanta pipeline (Takara Bio) and the Seurat R package (v3.1.5)[6] was used to perform further feature selection and clustering. Single cells with over 200 expressed genes were selected. We clustered the single cells by identifying the top 2,000 highly variable genes, computing principal component analysis based on the scaled expression values by total expression, and performing graph-based cluster detection using the top 10 principal components. Single cells were represented in a two-dimensional uniform manifold approximation and projection plane, and clusters were annotated according to the marker gene composition.

**Whole-genome/whole-exome sequencing analysis**

We performed whole-genome sequencing (WGS), whole-exome sequencing (WES), and SNP array experiments using peripheral blood mononuclear cells from the donor (termed control), the MCB, expansion cultures of the MCB, hiPSC-CMs, and hiPSC-CM patches.

We prepared 200 ng and 100 ng of genomic DNA as starting material for WGS and WES, respectively. Following the manufacturers' protocols, libraries for WGS were generated with the KAPA Hyper Prep Kit (Kapa Biosystems, Wilmington, MA) without PCR from fragmented genomic DNA sheared by Covaris LE220 (Covaris, Brighton, UK). For WES, adapter-ligated libraries were prepared with the KAPA Hyper Prep Kit (Kapa Biosystems) and sequencing libraries were constructed using the SeqCap EZ Human Exome Library v3.0 (Roche). Cluster generation was performed with the HiSeq

PE Cluster Kit v4 (Illumina) using Illumina cBot. Sequencing was performed using the HiSeq2500 platform in the 126 paired-end mode. After FASTQ files were generated (via bcl2fastq v2.17.1.14; Illumina) and adapter trimming was performed using cutadapt 1.10 [7], FASTQ files were mapped to the reference human genome (hg19 with decoy plasmid sequences for establishing hiPSCs and PhiX sequence) using BWA MEM (v0.7.15; with the default parameters, except for the use of the T-0 option for Genomon2) [8], and duplicated reads were removed using NovoSort (Novocraft; v1.03.09). The depths of coverage of WGS and WES data were 56X-83X and 84X-114X on average, respectively. To call single nucleotide variants (SNVs) or insertions/deletions (indels) in the test samples compared with the control sample, bam files were analysed via Genomon (v1.0.1)[9] and Genomon2 (v2.3.0) [10] with the EB call. The significance level, as per Fisher's exact test used in Genomon, was $P < 0.001$ for WGS and WES. In Genomon2, $P < 0.1$ was used for WGS and WES, and the significance levels of the EB call were set as $P < 0.001$ and $P < 0.0001$ for WGS and WES, respectively. After calling SNV/indels with Genomon and Genomon2, mutations whose variant allele frequencies were < 0.05 and fewer than five times those of the control sample were discarded. Thereafter, functional annotation was performed using ANNOVAR[11]; mutations were restricted to CDS and splicing sites. To extract potentially pathogenic mutations further, we excluded synonymous mutations and focused on mutations possibly related to cancer or other diseases. Mutations registered in the population databases, SNP131[12], esp6500si_all (> 0.01)[13], 1000g2014oct_all (> 0.01)[14], HGVD (v 2.0.0; > 0.01)[15], and 1KJPN (v1; > 0.01)[16] were discarded, but those registered in HGMD Pro (2016.4) [17], COSMIC79_position [18], and in the COSMIC Cancer Gene Census (v79)[19], or the Shibata's list [20] were retained. Mutations that passed these filters were reported. WGS data were also used to call CNVs with VarScan2 (v2.4.2) [21] in combination with the Otsu's threshold method [22] and Delly [23] (v0.7.3) by comparing the test samples and the control sample. Finally, we curated candidate CNVs manually based on overall trends in the depth of coverage, mapping status, and characteristics of the genomic regions, which were assessed by observing the positions of the candidate CNVs within the genome browser. We investigated genomic mutations by comparing the test samples and the control sample.

**Electron microscopy**

Cardiac tissues were prefixed with Karnovsky fixative containing 2.5% glutaraldehyde and 2% paraformaldehyde in 0.1 M cacodylate buffer (pH 7.4) for 2 h at 4 °C and postfixed in 2% osmium tetroxide (Nisshin EM, Tokyo, Japan) for 2 h at 4 °C. The samples were then immersed in 0.5% uranyl acetate (Fujifilm) for 3 h at room temperature, dehydrated in ethanol (50%, 70%, 80%, 90%, 95%, and 100%; Muto Pure Chemicals, Tokyo, Japan) and propylene oxide (Sigma-Aldrich), and embedded in epoxy resin. Semi-thin sections (0.5 μm) were stained with 0.1% toluidine blue (Merck, Darmstadt, Germany) solution and examined under a light microscope. Ultrathin sections were made with an EM UC7 ultra-microtome (Leica Microsystems, Wetzlar, Germany). These sections were counterstained with uranyl acetate and lead citrate before examination with an H-7500 electron microscope (Hitachi High-Technologies, Tokyo, Japan) at 80 kV.

**Intracellular calcium imaging of hiPSC-CMs**

The hiPSC-CMs were seeded onto 96-well plates at 1–2 × $10^5$ cells/well. Intracellular calcium imaging analysis was performed as previously described[24]. Cells were loaded with 2 μM Cal-520™ (AAT Bioquest, Sunnyvale, CA) in PBS for 2 h at 37 °C. Next, the loading buffer was replaced with culture medium. The basal activity was recorded with FDSS/μCELL (Hamamatsu Photonics, Shizuoka, Japan). Subsequently, 0.1, 1, 10, 100, 1,000 nM of isoproterenol (Merck) or E-4031 (Merck) was added to the culture medium, and cells were monitored for 30 min after the addition of each drug.

Parameters, such as the peak rate, peak-to-peak time, up-stroke slope, down-stroke slope, and 90% peak-width duration (PWD90) were calculated from the ratio of fluorescence intensity before and after drug administration to characterise the intracellular calcium levels. The data are expressed as the mean ±SD.

**Contraction properties of hiPSC-CMs**

hiPSC-CMs were seeded onto 96-well plates at 1–2 × $10^5$ cells/well. Cell motion analysis was performed as previously described[24]. The motion was recorded at a frame rate of 150 fps, a depth of 8 bits, and a resolution of 1024 × 1024 pixels with a Cell Motion Imaging System (SI8000; Sony Biotechnology, Tokyo, Japan). Subsequently, 0.1, 1, 10, 100, 1,000 nM of isoproterenol (Merck) or E-4031 (Merck) were added to the culture medium, and cells were monitored for 30 min after the addition of each drug. Data on arrhythmia-like abnormal contractions after the addition of E-4031 were excluded from the analysis as they cannot be compared with data on contractions at regular intervals. The relative change of each parameter, such as beating rate, peak interval, contraction/relaxation velocity, and contraction relaxation duration (CRD) after drug administration was calculated using the samples pre-drug addition as the control group. The data are expressed as the mean ±SD.

**Generation of the porcine chronic myocardial infarction model and hiPSC-CM patch transplantation**

A chronic myocardial infarction model was generated by placing an ameroid constrictor (COR-4.0-SS; Research Instruments SW, Escondido, CA) around the proximal left anterior descending coronary artery (LAD) and ligation of the distal LAD after the second diagonal branch[25] in healthy 7–8-month-old mature female Clawn minipigs (Kagoshima Miniature Swine Research Center, Kagoshima, Japan) weighing 20–25 kg. Four weeks after the procedure, we selected chronic heart failure models using cardiac echocardiography and cardiac MRI. The minipigs were randomly divided into two groups: those receiving hiPSC-CM patch transplantation (hiPSC-CM patch group; n = 7) or those sham-operated (sham group; n = 4). Transplantation of hiPSC-CM patches was performed under general anaesthesia; tacrolimus was also administered (5 mg) orally. All animals received immunosuppressive drugs, such as tacrolimus (0.75 mg/kg; Astellas Pharma Inc., Tokyo, Japan), methylprednisolone (20 mg; Takeda Pharmaceutical Co. Ltd, Osaka, Japan), and mycophenolate mofetil (500 mg; Teva Czech Industries sro., Opava, Czechia)[26], which were administered daily starting from 5 days before transplantation until euthanasia. The hiPSC-CM patches were transplanted via median sternotomy under general anaesthesia by inhalation of 2% isoflurane (Fujifilm) and continuous injection of 6 $mg \cdot kg^{-1}$ $h^{-1}$ propofol (Diprivan; AstraZeneca, Osaka, Japan).

In the hiPSC-CM patch group, two patches containing $1 \times 10^8$ cells were transplanted onto the infarcted myocardium. The minipigs were then allowed to recover in individual cages controlled at 20–22 °C. Later, the minipigs were humanely sacrificed for analysis according to the Osaka University Regulations on Animal Experiments.

**Cardiac echocardiography**

Transthoracic echocardiography was performed under general anaesthesia using a 5.0 MHz transducer (Aplio Artida; Toshiba Medical Systems, Tokyo, Japan). The left ventricular end-diastolic (LVEDV) and end-systolic (LVESV) volumes were calculated using the Teichholz formula[27]. The left ventricular ejection fraction (LVEF) was calculated as follows: LVEF (%) = 100 × (LVEDV − LVESV)/(LVEDV).

**Cardiac catheter**

Fluoroscopy-guided selective coronary angiography was performed by injecting iopamidol through a catheter (Vista Brite Tip; Cordis, Miami Lakes, FL) inserted from the right femoral artery in the supine position under general anaesthesia. A fluoroscopy-guided pressure wire (Radi Medical Systems, Uppsala, Sweden) was inserted to assess the myocardial microvascular resistance in the left circumflex coronary artery (LCx) (posterolateral and obtuse marginal branches) and right coronary artery (RCA) territories (Figure 5A), as described by Fearon et al.[28]. The coronary pressure wire was calibrated outside the body, equalised to the pressure reading from the guide catheter with the pressure sensor positioned at the ostium of the guide catheter, and then advanced to the distal two thirds of LCx and RCA. The index of microvascular resistance (IMR) was determined as follows: 3 mL of room temperature saline was injected into the cardiac catheter three times at rest, and the resting transit times were recorded and averaged. Maximal hyperaemia was then achieved using continuous intravenous adenosine at 180 $mg \cdot kg^{-1}$ $min^{-1}$ via a venous catheter. The maximal hyperaemic transit time was measured three times and averaged. The mean aortic and distal coronary pressures were recorded during the peak hyperaemia. An IMR $\leq 25$ was considered normal[29].

**Cardiac MRI**

Cardiac MRI was performed under general anaesthesia using a 1.5-T MR scanner (SIGNA EXCITE XI TwinSpeed; GE Medical Systems, Milwaukee, WI), just before and 12 weeks after transplantation. The images were then analysed using a commercial feature tracking software (2D CPA MR; Tom-Tec Imaging Systems, Unterschleissheim, Germany), a vector-based analysis tool based on a hierarchical algorithm that has been previously validated in clinical studies[30]. For each of the three short-axis plane cine images, the left ventricle endocardial border at the end-diastolic frame was manually drawn on a single frame by an expert reader. The software then automatically propagated the contour and followed its features throughout the cardiac cycle to draw the circumferential strain of the 17 segments, as per the American Heart Association model. The 17 segments were compiled into three territories, according to the coronary artery domination: LAD (territories 1, 2, 7, 8, 13, and 14), LCx (territories 5, 6, 11, 12, and 16), and RCA (territories 3, 4, 9, 10, and 15).

**Telemetered Holter electrocardiography**

To assess safety concerning arrhythmias, we recorded 24-h electrocardiograms using a Holter recorder (PhysioTel Digital, DSI, USA) 7 days before and on days 0–3, 7, 14, 28, 42, 56, 70, and 84 after implantation. The electrodes were placed in the chest and connected to the Holter recorder. The radiofrequency signals were monitored and saved on a personal computer near the cages using PhysioTel Digital System (DSI, USA.). All data were analysed using HEM data analysis software (Notocord Systems SAS, Croissy-sur-Seine, France).

**Soft agar colony formation assay**

Modified Eagle's Medium (Thermo Fisher Scientific) supplemented with 10% FBS and Bacto Agar (BD Biosciences) solution was added to a 60 mm dish and allowed to solidify. hiPSCs-CM patches were dissociated into single cells via treatment with 0.25% trypsin-EDTA solution (Thermo Fisher Scientific) and suspended in DMEM supplemented with 10% FBS. HeLa cells (JCRB Cell Bank, NIBIOHN, Osaka, Japan) were used as the positive control, and MRC-5 cells (DS Pharma Biomedical, Osaka, Japan) were used as the negative control. Cells were mixed with noble agar and spread onto the 60 mm dish pre-covered with a bottom layer. After placement onto the bottom agar layer, the top agar layers immediately solidified. The dishes were incubated with culture medium containing 10% FBS for three weeks at 37 °C and 5% $CO_2$. At the end of the incubation period, colonies were visualised via staining with ρ-iodonitrotetrazolium violet (Nacalai Tesque) for 6 h at 37 °C and 5% $CO_2$. Colony formation images were acquired using the inverted microscope IX73 (Olympus).

**General toxicity tests**

The hiPSC-CM patches were tested for general toxicity using immunodeficient NOD/Shi-scid, IL-2R γnull mice (NOG mice; In-Vivo Science Inc., Tokyo, Japan), which are further described in Table S7. Mice were housed with bedding (autoclaved-wood shavings; Rettenmaier Japan Co., Ltd., Tokyo, Japan) and provided water (tap water containing 2 ppm sodium hypochlorite) and food (Charles River Laboratories Japan, Inc., Kanagawa, Japan) *ad libitum*. One hiPSC-CM patch consisting of 1.9 million hiPSC-CMs was directly transplanted onto the surface of the left anterior wall of the heart. The mice were divided into six groups: (1) hiPSC-CM-receiving males; (2) hiPSC-CM-receiving females; (3) sham-operated (open chest) males; (4) sham-operated females; (5) non-operated males; and (6) non-operated females (n = 10 each). Twenty-eight days after transplantation, mice were euthanised by exsanguination under inhalation anaesthesia with isoflurane (Mylan Inc., Canonsburg, PA) and dissected. Gross abnormalities and the weight of the major organs were recorded. Peripheral blood was collected to conduct haematological and biochemical evaluations.

**Tumourigenicity assay**

The hiPSC-CM patch was tested for tumourigenicity using immunodeficient NOG mice. As described in the above section, one hiPSC-CM patch consisting of 1.9 million hiPSC-CMs was directly transplanted onto the surface of the left anterior wall of the heart (n = 10 in each group). Mouse survival and body weight following transplantation were recorded. Mice were euthanised and dissected 16 weeks after transplantation. The major organs and tissues were carefully observed, and any gross pathological findings were collected and stored for further examination.

**Histological analysis**

All autopsy tissue specimens of murine and porcine hearts transplanted with hiPSC-CMs were fixed in 10% buffered formalin (Fujifilm) and embedded in paraffin using a Microm STP 120 Spin Tissue Processor (STP120-3; Thermo Fisher Scientific). Serial paraffin-embedded sections were cut at a thickness of 0.5 μm using a Microm HM 430 system (MIC 990010; Thermo Fisher Scientific), deparaffinised in xylene (Fujifilm), dehydrated in a graded series of ethanol (Fujifilm), and stained with haematoxylin and eosin (H&E; Muto Pure Chemicals). The sections were then imaged using a light microscope (DM4000B; Leica). For analysing fibrosis, paraffin-embedded sections were stained with Picrosirius red (Fujifilm) and imaged under a microscope. The percentage of the fibrotic area in the entire tissue was measured using MetaMorph software for Windows (Universal Imaging Corporation, Downingtown, PA). Immunostaining was performed using anti-Ki-67 (1:100; M7240; Dako, Glostrup, Denmark) and anti-lamin antibodies (1:250; ab108595; Abcam). Briefly, deparaffinised, dehydrated tissue sections were processed for antigen retrieval via autoclaving in 0.01 M citrate buffer (Dako). The sections were immersed in methanol (Fujifilm) containing 3% hydrogen peroxide (Fujifilm) and the slides were incubated overnight at 4 °C with the indicated primary antibodies. Subsequently, the sections were incubated with a biotinylated anti-mouse IgG antibody (K0675; Dako), further incubated with peroxidase-conjugated streptavidin (Dako), and then visualised using the biphenyl-3,30,4,40-tetramine (DAB) solution (Fujifilm). The sections were imaged under a light microscope (Leica).

**Fluorescent *in situ* hybridisation**

hiPSC-CMs at 1 week after transplantation were assessed via fluorescent *in situ* hybridisation (FISH) using a human-specific genomic probe labelled as previously described[31]. Briefly, 3 mm sections were deparaffinised, washed in phosphate-buffered saline for 5 min, digested in pepsin solution (0.1% in 0.1 N HCl) at 37 °C for 10 min, and dehydrated. A human-specific FISH probe (Chromosome Science Labo Inc., Sapporo, Japan) labelled with Cy3 was applied to the pre-treated sections, which were covered by cover slips and simultaneously denatured at 90 °C for 10 min. Hybridisation was carried out at 37 °C overnight. Sections were then washed with 50% formamide, 2X SSC at 37 °C for 20 min, and 1X SSC for 15 min at room temperature, followed by counterstaining with 4′,6-diamidino-2-phenylindole (DAPI) and mounting.

#### Supplementary References

1. Okita K, Yamakawa T, Matsumura Y, Sato Y, Amano N, Watanabe A, Goshima N, Yamanaka S. An efficient nonviral method to generate integration-free human-induced pluripotent stem cells from cord blood and peripheral blood cells. Stem Cells. 2013;31(3):458-66.

2. Nakagawa M, Taniguchi Y, Senda S, Takizawa N, Ichisaka T, Asano K, Morizane A, Doi D, Takahashi J, Nishizawa M, Yoshida Y, Toyoda T, Osafune K, Sekiguchi K, Yamanaka S. A novel efficient feeder-free culture system for the derivation of human induced pluripotent stem cells. Sci Rep 2014;4:3594.

3. Ito E, Miyagawa S, Takeda M, Kawamura A, Harada A, Iseoka H, Yajima S, Sougawa N, Mochizuki-Oda N, Yasuda S, Sato Y, Sawa Y. Tumorigenicity assay essential for facilitating safety studies of hiPSC-derived cardiomyocytes for clinical application. Sci Rep 2019;9(1):1881.

4. Ito E, Miyagawa S, Yoshinori Yoshida, Sawa Y. Efficient method to dissociate induced pluripotent stem cell-derived cardiomyocyte aggregates into single cells. Methods Mol Biol. 2021;2320.

5. Sougawa N, Miyagawa S, Fukushima S, Kawamura A, Yokoyama J, Ito E, Harada A, Okimoto K, Mochizuki-Oda N, Saito A, Sawa Y. Immunologic targeting of CD30 eliminates tumourigenic human pluripotent stem cells, allowing safer clinical application of hiPSC-based cell therapy. Sci Rep 2018;8(1):3726.

6. Butler A, Hoffman P, Smibert P Papalexi E, Satija R. Integrating single-cell transcriptomic data across different conditions, technologies, and species. Nat Biotechnol 2018;36(5):411–420.

7. Martin M. Cutadapt removes adapter sequences from high-throughput sequencing reads. EMBnet J 2011;17(1):10–12.

8. Li H, Durbin R. Fast and accurate short read alignment with Burrows-Wheeler transform. Bioinformatics 2009;25(14):1754–1760.

9. Yoshida K, Sanada M, Shiraishi Y, Nowak D, Nagata Y, Yamamoto R, Sato Y, Sato-Otsubo A, Kon A, Nagasaki M, Chalkidis G. Frequent pathway mutations of splicing machinery in myelodysplasia. Nature 2011;478(7367):64–69.

10. Shiraishi Y, Sato Y, Chiba K, Okuno Y, Nagata Y, Yoshida K, Shiba N, Hayashi Y, Kume H, Homma Y, Sanada M, Ogawa S, Miyano S. An empirical

Bayesian framework for somatic mutation detection from cancer genome sequencing data. Nucleic Acids Res 2013;41(7):e89.

11. Wang K, Li M, Hakonarson H. ANNOVAR: functional annotation of genetic variants from high-throughput sequencing data. Nucleic Acids Res 2010;38(16):e164.

12. Sherry ST, Ward MH, Kholodov M, et al. dbSNP: the NCBI database of genetic variation. Nucleic Acids Res 2001;29(1):308–311.

13. Exome Variant Server. NHLBI GO Exome Sequencing Project (ESP), Seattle, WA. http://evs.gs.washington.edu/EVS/ (July 2016)

14. Auton A, Brooks LD, Durbin RM, Garrison EP, Kang HM, Korbel JO, Marchini JL, McCarthy S, McVean GA, Abecasis GR. A global reference for human genetic variation. Nature 2015;526(7571):68–74.

15. Higasa K, Miyake N, Yoshimura J, Okamura K, Niihori T, Saitsu H, Doi K, Shimizu M, Nakabayashi K, Aoki Y, Tsurusaki Y, Morishita S, Kawaguchi T, Migita O, Nakayama K, Nakashima M, Mitsui J, Narahara M, Hayashi K, Funayama R, Yamaguchi D, Ishiura H, Ko WY, Hata K, Nagashima T, Yamada R,

Matsubara Y, Umezawa A, Tsuji S, Matsumoto N, Matsuda F. Human genetic variation database, a reference database of genetic variations in the Japanese population. Journal Hum Genet 2016;61(6):547–553.

16. Nagasaki M, Yasuda J, Katsuoka F, Nariai N, Kojima K, Kawai Y, Yamaguchi-Kabata Y, Yokozawa J, Danjoh I, Saito S, Sato Y, Mimori T, Tsuda K, Saito R, Pan X, Nishikawa S, Ito S, Kuroki Y, Tanabe O, Fuse N, Kuriyama S, Kiyomoto H, Hozawa A, Minegishi N, Douglas Engel J, Kinoshita K, Kure S, Yaegashi N; ToMMo Japanese Reference Panel Project, Yamamoto M. Rare variant discovery by deep whole-genome sequencing of 1,070 Japanese individuals. Nat Commun 2015;6:8018.

17. Stenson PD, Mort M, Ball EV, Evans K, Hayden M, Heywood S, Hussain M, Phillips AD, Cooper DN. The Human Gene Mutation Database: towards a comprehensive repository of inherited mutation data for medical research, genetic diagnosis and next-generation sequencing studies. Hum Genet 2017;136(6):665–677.

18. Tate JG, Bamford S, Jubb HC, Sondka Z, Beare DM, Bindal N, Boutselakis H, Cole CG, Creatore C, Dawson E, Fish P, Harsha B, Hathaway C, Jupe SC, Kok

CY, Noble K, Ponting L, Ramshaw CC, Rye CE, Speedy HE, Stefancsik R, Thompson SL, Wang S, Ward S, Campbell PJ, Forbes SA. COSMIC: the Catalogue Of Somatic Mutations In Cancer. Nucleic Acids Res 2019;47(D1):D941–D947.

19. Sondka Z, Bamford S, Cole CG, Ward SA, Dunham I, Forbes SA. The COSMIC Cancer Gene Census: describing genetic dysfunction across all human cancers. Nat Rev Cancer 2018;18(11):696–705.

20. Nakahata T, Okano H. Current perspective on evaluation of tumorigenicity of cellular and tissue-based products derived from induced pluripotent stem cells (iPSCs) and iPSCs as their starting materials (provisional translation). Pharmaceuticals and Medical Devices Agency. http://www.pmda.go.jp/files/000152599.pdf

21. Koboldt DC, Zhang Q, Larson DE, Shen D, McLellan MD, Lin L, Miller CA, Mardis ER, Ding L, Wilson RK. VarScan 2: somatic mutation and copy number alteration discovery in cancer by exome sequencing. Genome Res 2012;22(3):568–576.

22. Otsu N. A Threshold Selection Method from Gray-Level Histograms. IEEE Trans Syst Man Cybern 1979;9(1):62–66.

23. Rausch T, Zichner T, Schlattl A, Stütz AM, Benes V, Korbel JO. DELLY: structural variant discovery by integrated paired-end and split-read analysis. Bioinformatics 2012;28(18):i333-i9.

24. Takeda M, Miyagawa S, Fukushima S, Saito A, Ito E, Harada A, Matsuura R, Iseoka H, Sougawa N, Mochizuki-Oda N, Matsusaki M. Development of In Vitro Drug-Induced Cardiotoxicity Assay by Using Three-Dimensional Cardiac Tissues Derived from Human Induced Pluripotent Stem Cells. Tissue Eng Part C Methods 2018;24(1):56–67.

25. Teramoto N, Koshino K, Yokoyama I, Miyagawa S, Zeniya T, Hirano Y, Fukuda H, Enmi J, Sawa Y, Knuuti J, Iida H. Experimental pig model of old myocardial infarction with long survival leading to chronic left ventricular dysfunction and remodeling as evaluated by PET. J Nucl Med 2011;52(5):761–768.

26. Kawamura M, Miyagawa S, Fukushima S, Saito A, Miki K, Ito E, Sougawa N, Kawamura T, Daimon T, Shimizu T, Okano T, Toda K, Sawa Y. Enhanced

survival of transplanted human induced pluripotent stem cell-derived cardiomyocytes by the combination of cell sheets with the pedicled omental flap technique in a porcine heart. Circulation 2013;128(11 Suppl 1):S87–S94.

27. Teichholz LE, Kreulen T, Herman MV, Gorlin R. Problems in echocardiographic volume determinations: Echocardiographic-angiographic correlations in the presence or absence of asynergy. Am J Cardiol 1976;37(1):7–11.

28. Fearon WF, Balsam LB, Farouque HM, Caffarelli AD, Robbins RC, Fitzgerald PJ, Yock PG, Yeung AC. Novel index for invasively assessing the coronary microcirculation. Circulation 2003;107(25):3129–3132.

29. Melikian N, Vercauteren S, Fearon WF, Cuisset T, MacCarthy PA, Davidavicius G, Aarnoudse W, Bartunek J, Vanderheyden M, Wyffels E, Wijns W, Heyndrickx GR, Pijls NH, de Bruyne B. Quantitative assessment of coronary microvascular function in patients with and without epicardial atherosclerosis. EuroIntervention 2010;5(8):939–945.

30. Obokata M, Nagata Y, Wu VC, Kado Y, Kurabayashi M, Otsuji Y, Takeuchi M. Direct comparison of cardiac magnetic resonance feature tracking and 2D/3D

echocardiography speckle tracking for evaluation of global left ventricular strain. Eur Heart J Cardiovasc Imaging 2016;17(5):525–532.

31. Kawamura M, Miyagawa S, Miki K, Saito A, Fukushima S, Higuchi T, Kawamura T, Kuratani T, Daimon T, Shimizu T, Okano T, Sawa Y. Feasibility, safety, and therapeutic efficacy of human induced pluripotent stem cell-derived cardiomyocyte sheets in a porcine ischaemic cardiomyopathy model. Circulation 2012;126(11 Suppl 1):S29–S37.

#### Supplementary Figure Legends

**Figure S1. Quality check of hiPSCs, hiPSC-CMs, and the hiPSC-CM patch.**

**Figure S2. Characterisation of hiPSC-CMs.**

A: Number of cells per 100 mL bioreactor during cardiomyogenic differentiation-induction. The cell number was measured 4, 8, 10, 12, 14, 16, and 25 days after induction.

B: Representative immunofluorescence staining images of differentiated embryoid bodies during cardiomyogenic differentiation-induction. Upper panel, staining of Lin28A (red), a marker of undifferentiated stem cells, and cardiac troponin T (cTNT; green), a cardiomyocyte marker; nuclei (blue, Hoechst). Lower panel, staining of α-actinin (red) and cTNT (green), both cardiomyocyte markers; nuclei (blue, Hoechst). Scale bar: 20 μm.

C: Gene expression analysis in the context of the cardiac differentiation-induction process.

qPCR of markers of (C-1) pluripotency: *POU5F1*, *SOX2*, *NANOG*, and *Lin28A*; (C-2) early mesoderm: *MESP1*, Brachyury, and *EOMES*; (C-3) cardiac progenitor cells: *Isl1*, *PDGFRA*, and *MEF2C*; and (C-4) cardiomyocytes: *TNNT2*, *ACTN2*, and *MYH7*. Data shown are of one representative experiment for each marker, and are normalised to peak expression ± SD.

**Figure S3. Gene expression pattern of hiPSC-CM.**

A: Heat map of normalised profiling data comparing undifferentiated hiPSCs and hiPSC-CMs. Expression of these 84 genes represents the stem cell pathway.

B: Heat map of normalised profiling data comparing undifferentiated hiPSCs, hiPSC-CMs, human foetal hearts, and human adult hearts. These 23 genes represent the cardiac differentiation-associated genes.

C: Relative comparison of gene expression associated with ion channels in hiPSCs, hiPSC-CMs, and the adult heart.

**Figure S4. Intracellular calcium levels of hiPSC-CM following drug administration.**

A: Schematic diagram of the calcium transient analysis.

B, C: Representative calcium transient waveform after the addition of isoproterenol (B) or E-4031 (C) and quantitative analysis of the changes in calcium levels after drug administration. The relative change in each parameter, such as the peak rate, peak-to-peak time, up-stroke slope, down-stroke slope, and 90% peak-width duration

(PWD90) after drug administration, was calculated using the samples pre-drug addition as the control group. Data are presented as the mean ± SD. *$P$ < 0.05, **$P$ < 0.01 vs. vehicle control group.

**Figure S5. Contraction properties of hiPSC-CM following drug administration.**

A: Schematic diagram of the contraction analysis.

B, C: Representative contraction waveform after the addition of isoproterenol (B) or E-4031 (C) and quantitative analysis of the changes in contraction parameters after drug administration. The relative change in each parameter, such as beating rate, peak interval, contraction/relaxation velocity, and contraction relaxation duration (CRD) after drug administration, was calculated using the samples pre-drug addition as the control group. Data are presented as the mean ± SD. *$P$ < 0.05, **$P$ < 0.01 vs. vehicle control.

**Figure S6. Characterisation of the hiPSC-CM patch.**

A: Transmission electronic microscopy of the hiPSC-CM patch ultrastructural features. Z, Z-line; Mt, mitochondria; R, ribosomes; N, nucleus. Scale bar = 2 μm (magnification, 3,000×).

B: Expression of angiogenic cytokines under normoxic or hypoxic conditions.

Data are presented as the mean ± SD. *$P < 0.01$.

**Figure S7. Experimental protocol for determining efficacy using the porcine MI model.**

**Figure S8. Post-transplant hiPSC-CM status and effect on organs.**

A: Heart; B: Lung; C: Liver; D: Kidney; E: Spleen, and corresponding H&E-stained images. Scale bar: 200 μm.

#### Supplementary Tables

##### Table S1. List of antibodies used in this study

| Antibody | Company | Catalogue No. |
| --- | --- | --- |
| cTNT | Abcam | ab45932 |
| cTNT | Thermo Fisher Scientific | MS-295-P |
| cTNT | Santa Cruz Biotechnology | sc-20025 |
| α-Actinin | Sigma | A7811 |
| α-Actinin | Abcam | ab68167 |
| Connexin43 | Sigma | C6219 |
| MLC2v | Proteintech | 10906-1-AP |
| MLC2a | Synaptic Systems | 311 011 |
| α-MHC | Sigma | HPA001349 |
| α-MHC | R&D | 940344 |
| β-MHC | Sigma | M8421 |
| LIN28A | LSBio | LS-B5073 |
| CD31 | Abcam | ab28364 |
| αSMA | DAKO | M0851 |

| | | |
|---|---|---|
| αSMA | Abcam | ab32575 |
| N-cadherin | Abcam | ab18203 |
| Vimentin | Abcam | ab92547 |
| Collagen I | Abcam | ab34710 |
| Laminin | Sigma | L9393 |
| Alexa Fluor 488 goat anti-mouse | Thermo Fisher Scientific | A11001 |
| Alexa Fluor 488 donkey anti-mouse | Thermo Fisher Scientific | A21202 |
| Alexa Fluor 555 donkey anti-mouse | Thermo Fisher Scientific | A31570 |
| Alexa Fluor 555 goat anti-rabbit | Thermo Fisher Scientific | A21428 |
| Alexa Fluor 488 donkey anti-rabbit | Thermo Fisher Scientific | A21206 |
| Alexa Fluor 555 donkey anti-rabbit | Thermo Fisher Scientific | A31572 |

cTNT, cardiac troponin T; MLC2a, atrial isoform of the myosin light chain 2; MLCv, ventricular isoform of myosin light chain; MHC, myosin heavy chain; SMA, smooth muscle actin.

**Table S2. List of plasmids used for hiPSC reprogramming**

| Plasmid name | Gene | Source or reference |
|---|---|---|
| pCE-hSK | *SOX2, KLF4* | [1] |
| pCE-hUL | *L-MYC, LIN28* | [1] |
| pCE-hOCT3/4 | *OCT3/4* | [1] |
| pCE-mp53DD | *Trp53* | [1] |
| pCXB-EBNA1 | *EBNA1* | [1] |

**Table S3. Characterisation of hiPSCs**

| Assay | Method | Criteria | Results |
|---|---|---|---|
| Morphology | Microscopic examination | Human ES cell-like | Human ES cell-like |
| Remaining plasmid vector | qPCR | Not detected | Not detected |
| Karyotype analysis | Conventional Giemsa G-band | Normal (22 pairs of autosomal chromosomes and one pair of sex chromosomes) | Normal (22 pairs of autosomal chromosomes and one pair of sex chromosomes) |
| Expression of pluripotent markers | RNA microarray | *POU5F1*: ≥ 4%, *NANOG*: ≥ 5%, vs. *GAPDH* | Within the standard values |
| | Flow cytometry | *SSEA4*: ≥ 90%, *TRA-1-60*: ≥ 90%, *TRA-2-49*: ≥ 90% | Within the standard values |

| | | | |
|---|---|---|---|
| Expression of differentiation resistance markers | RNA microarray | *C4orf51:* ≤ 0.04%, *ABHD12B:* ≤ 0.07%, *HHLA1:* ≤ 0.25% vs. *GAPDH* | Within the standard values |
| Doubling time | Calculated from cell growth curve | 15–45 h | Within the standard values |
| Sterility testing | BacT/ALERT ® MB | Negative | Negative |
| Mycoplasma testing | PCR | Negative | Negative |
| Endotoxin testing | Kinetic-turbidimetric technique | ≤ 5 EU/mL | Within the criterion |
| Viral testing | PCR (*HBC, HCV, HIV, HTLV*, and *Parvovirus 19*) | All negative | All negative |
| HLA typing | PCR-SBT (one of each from *HLA-A*, *HLA-B*, and *HLA-DR*) | Match with the donor blood cell profile | Match with the donor cell profile |

| | | | |
|---|---|---|---|
| STR genotyping | PCR-capillary electrophoresis | Match with the donor blood cell profile | Match with the donor cell profile |

ES, embryonic stem cell; qPCR, quantitative polymerase chain reaction; HLA, human leukocyte antigen; STR, short tandem repeat.

**Table S4. Quality test of hiPSC-CMs**

| Assay | Method | Criteria | Results |
|---|---|---|---|
| Viability | Trypan Blue exclusion test | ≥ 40% | 65.7% |
| Purity of cardiomyocytes | Flow cytometry | cTNT-positive rate ≥ 50% | 73.4% |
| Sterility testing | Membrane filtration method | Negative | Negative |
| Mycoplasma testing | Nested PCR | Negative | Negative |
| Endotoxin testing | Turbidimetric technique | < 1.0 EU/mL | < 0.194 EU/mL |

hiPSC-CM, human induced pluripotent stem cell-derived cardiomyocyte; cTNT, cardiac troponin T

**Table S6. Representative results of telemetered Holter electrocardiography**

| ID | Group | Pre operation | After transplantation | | | | | | | |
|---|---|---|---|---|---|---|---|---|---|---|
| | | | 0–72 h | 1 w | 2 w | 4 w | 6 w | 8 w | 10 w | 12 w |
| 1 | Sham | 0 | 0 | 0 | 0 | 0 | 0 | 0 | 0 | 0 |
| 2 | Sham | 0 | 0 | 0 | 0 | 0 | 0 | 0 | 0 | 0 |
| 3 | Sham | 0 | 0 | 0 | 0 | 0 | 0 | 0 | 0 | 0 |
| 4 | Sham | 0 | 0 | 0 | 0 | 0 | 0 | 0 | 0 | 0 |
| 5 | hiPSC-CMs | 0 | 0 | 0 | 0 | 0 | 0 | 0 | 0 | 0 |
| 6 | hiPSC-CMs | 0 | 0 | 0 | 0 | 0 | 0 | 0 | 0 | 0 |
| 7 | hiPSC-CMs | 0 | 0 | 0 | 0 | 0 | 0 | 0 | 0 | 0 |
| 8 | hiPSC-CMs | 0 | 0 | 0 | 0 | 0 | 0 | 0 | 0 | 0 |
| 9 | hiPSC-CMs | 0 | 0 | 0 | 0 | 0 | 0 | 0 | 0 | 0 |
| 10 | hiPSC-CMs | 0 | 0 | 0 | 0 | 0 | 0 | 0 | 0 | 0 |
| 11 | hiPSC-CMs | 0 | 0 | 0 | 0 | 0 | 0 | 0 | 0 | 0 |

w, weeks.

**Table S7. Summary of the general toxicity and tumourigenicity tests**

| | | General toxicity | Tumorigenicity |
|---|---|---|---|
| Recipient | Strain | NOD/Shi-scid, IL2RγKO Jic mouse | |
| | Age | 6–9 weeks old | |
| | Sex | Male, female | Female |
| | Group | 1) hiPSC-CM patch-receiving<br>2) sham-operated<br>3) non-operated | hiPSC-CM patch-receiving |
| | Number | n = 10 | n = 10 |
| Transplant | | $1.9 \times 10^6$ hiPSC-CMs per patch, 1 sheet per mouse | |
| Follow-up | | 28 days | 16 weeks |
| Assessment | Haematology | RBC, HGB, HCT, MCV, MCH, MCHC, PLT, WBC, Neut, Lympho, Mono, EO, and BASO | None |
| | Biochemistry | ALP, AST, ALT, TCHO, TG, BUN, and CRE | None |

| | | | |
|---|---|---|---|
| | Autopsy | Brain, spinal cord, pituitary, oculars, Harderian glands, tongue, salivary glands, thyroid glands, trachea, heart, lung, oesophagus, aorta, liver, gallbladder, stomach, small intestine, large intestine, spleen, pancreas, kidneys, adrenal glands, urinary bladder, sciatic nerve, muscle, skin, sternum, femur, ovary, uterus, vagina, testis*, epididymis*, vesicular glands*, prostate glands*, and cervical lymph node. | |
| | Wet weight | Brain, liver, spleen, kidneys, adrenal glands, ovary, uterus, testis*, vesicular glands*, and prostate glands* | Brain, liver, spleen, kidneys, pituitary, heart, and lung |
| | Pathology | H&E staining | H&E staining and immunohistochemistry as needed (anti-laminin, anti-Ki-67) |

*For male mice only. RBC, red blood cell; HGB, haemoglobin; HCT, haematocrit; MCV, mean corpuscular volume; MCH, mean corpuscular haemoglobin; MCHC, mean corpuscular haemoglobin concentration; PLT, platelet; WBC, white blood cell; Neut, neutrophils; Lympho, lymphocytes; Mono, monocytes; EO, eosinophil; BASO, basophil; ALP, alkaline phosphatase; AST, aspartate aminotransferase; ALT, alanine aminotransferase; TCHO, total cholesterol; TG, triglyceride; BUN, blood urea nitrogen; CRE, creatinine; H&E, haematoxylin and eosin.

**Table S8. List of primers used in this study**

| Gene Symbol | Primer | |
|---|---|---|
| **Human** | | |
| *GAPDH* | SYBR | F: 3′- CAATGACCCCTTCATTGACC-5′<br>R: 5′- TTGATTTTGGAGGGATCTCG-3′ |
| *POU5F1* | SYBR | F: 3′-GAAACCCACACTGCAGCAGA-5′<br>R: 5′-TCGCTTGCCCTTCTGGCG-3′ |
| *SOX2* | SYBR | F: 3′-GCGCCCTGCAGTACAACTC -5′<br>R: 5′-CGGACTTGACCACCGAACC-3′ |
| *NANOG* | SYBR | F: 3′-CTCAGCTACAAACAGGTGAAGAC-5′<br>R: 5′-TCCCTGGTGGTAGGAAGAGTAAA-3′ |
| *Lin28A* | SYBR | F: 3′-CACGGTGCGGGCATCTG-5′<br>R: 5′-CCTTCCATGTGCAGCTTACTC-3′ |
| *MESP1* | SYBR | F: 3′-CAACTGACGCCGTCTCTGTGA-5′<br>R: 5′-GTCTGCCAAGGAACCACTTCG-3′ |
| *Brachyury* | SYBR | F: 3′- AATTGGTCCAGCCTTGGAAT-5′<br>R: 5′- CGTTGCTCACAGACCACA-3′ |
| *EOMES* | SYBR | F: 3′-CTTGCTAGGCCTCTGCTGTGTG-5′<br>R: 5′-TTGGTGACTCCTTAGCTTGCTCTCT-3′ |
| *Islet1* | SYBR | F: 3′-TTTATTGTCGGAAGACTTGCCACTT -5′<br>R: 5′-TCAAAGACCACCGTACAACCTTTATCT-3′ |
| *PDGFRA* | SYBR | F: 3′-TTGCTGTGAGCCTTGCATGA-5′<br>R: 5′-GTGGGAGCATTTGTTAGGACTGG-3′ |
| *MEF2C* | SYBR | F: 3′- TCGCTTGTAAATGAGGGCATACAA -5′<br>R: 5′- GTCCAGCTTATGCCGCTGTG-3′ |
| *TNNT2* | SYBR | F: 3′-GGCAGCTCCTGTTTGGAAATG-5′<br>R: 5′-TTATTACTGGTGTGGAGTGGGTGTG-3′ |
| *ACTN2* | SYBR | F: 3′-TTTCCCTGTGTGTTGGTTGC -5′<br>R: 5′-TGATTACACTCCGCACATTTCA-3′ |
| *MYH6* | SYBR | F: 3′-GAGATTTCTCCAACCCAG-5′<br>R: 5′-CCAGGGTGATGGAGAAGGAG-3′ |
| *MYH7* | SYBR | F: 3′-TTTCCCTGTGTGTTGGTTGC -5′<br>R: 5′-TGATTACACTCCGCACATTTCA-3′ |

| | | |
|---|---|---|
| *KCNQ1* | SYBR | F: 3′-GAGATTTCTCCAACCCAG-5′<br>R: 5′-CCAGGGTGATGGAGAAGGAG-3′ |
| *KCNH2* | SYBR | F: 3′-TTTCCCTGTGTGTTGGTTGC -5′<br>R: 5′-TGATTACACTCCGCACATTTCA-3′ |
| *CACNA1C* | SYBR | F: 3′-GAGATTTCTCCAACCCAG-5′<br>R: 5′-CCAGGGTGATGGAGAAGGAG-3′ |
| *SCN5A* | SYBR | F: 3′-TTTCCCTGTGTGTTGGTTGC -5′<br>R: 5′-TGATTACACTCCGCACATTTCA-3′ |
| *SERCA2* | SYBR | F: 3′-GAGATTTCTCCAACCCAG-5′<br>R: 5′-CCAGGGTGATGGAGAAGGAG-3′ |
| **Porcine** | | |
| *GAPDH* | TaqMan | Ss03374854_g1 |
| *SDF-1* | TaqMan | Ss03391855_m1 |
| *VEGF* | TaqMan | Ss03393993_m1 |
| *basic FGF* | TaqMan | Ss03375809_u1 |
| *HGF* | TaqMan | AJVI4PJ |

F, forward; R, reverse

#### Supplementary Figures

**Figure S1. Quality check of hiPSC, hiPSC-CMs, and the hiPSC-CM patch**

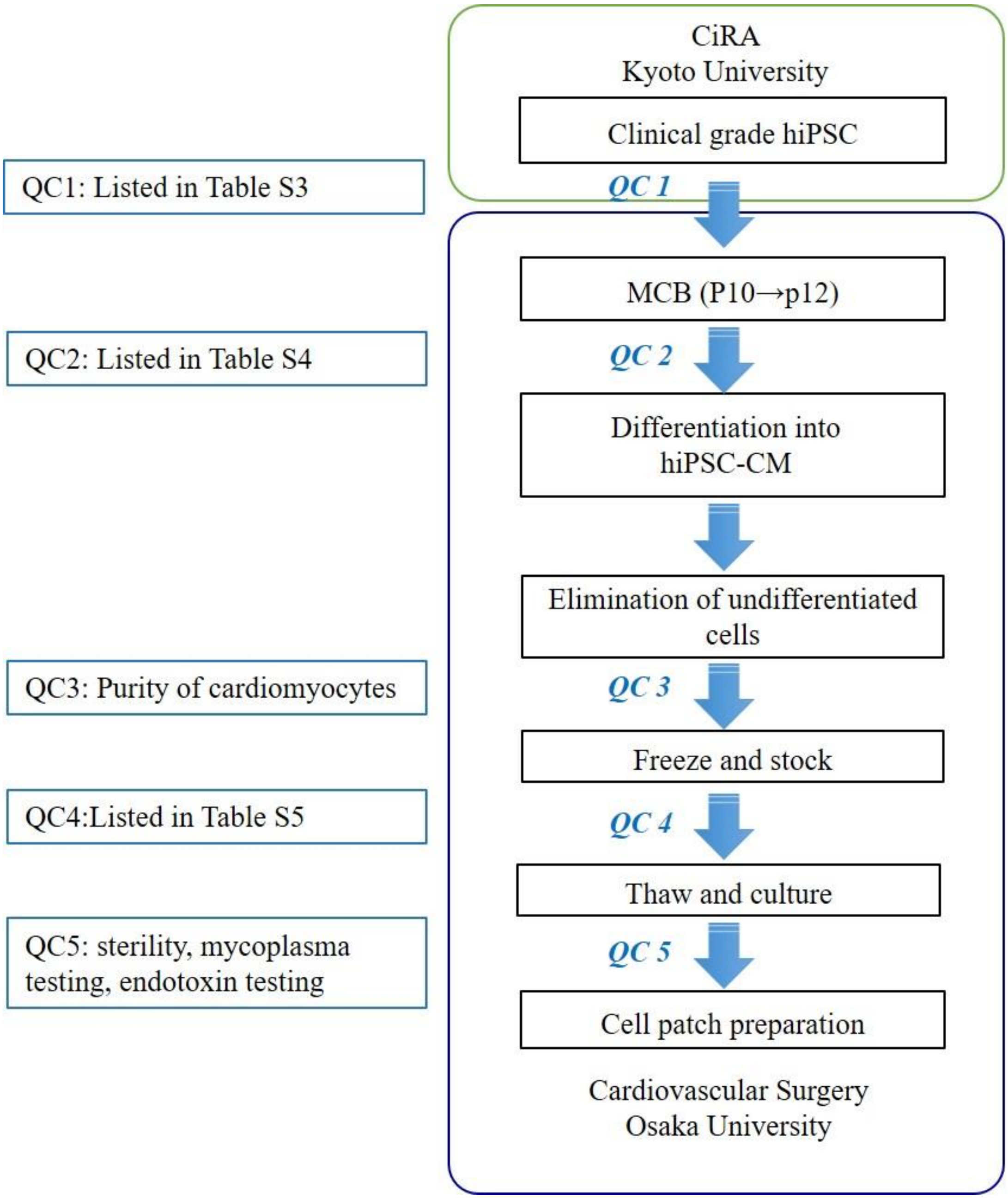

**Figure S2. Characterisation of hiPSC-CMs**

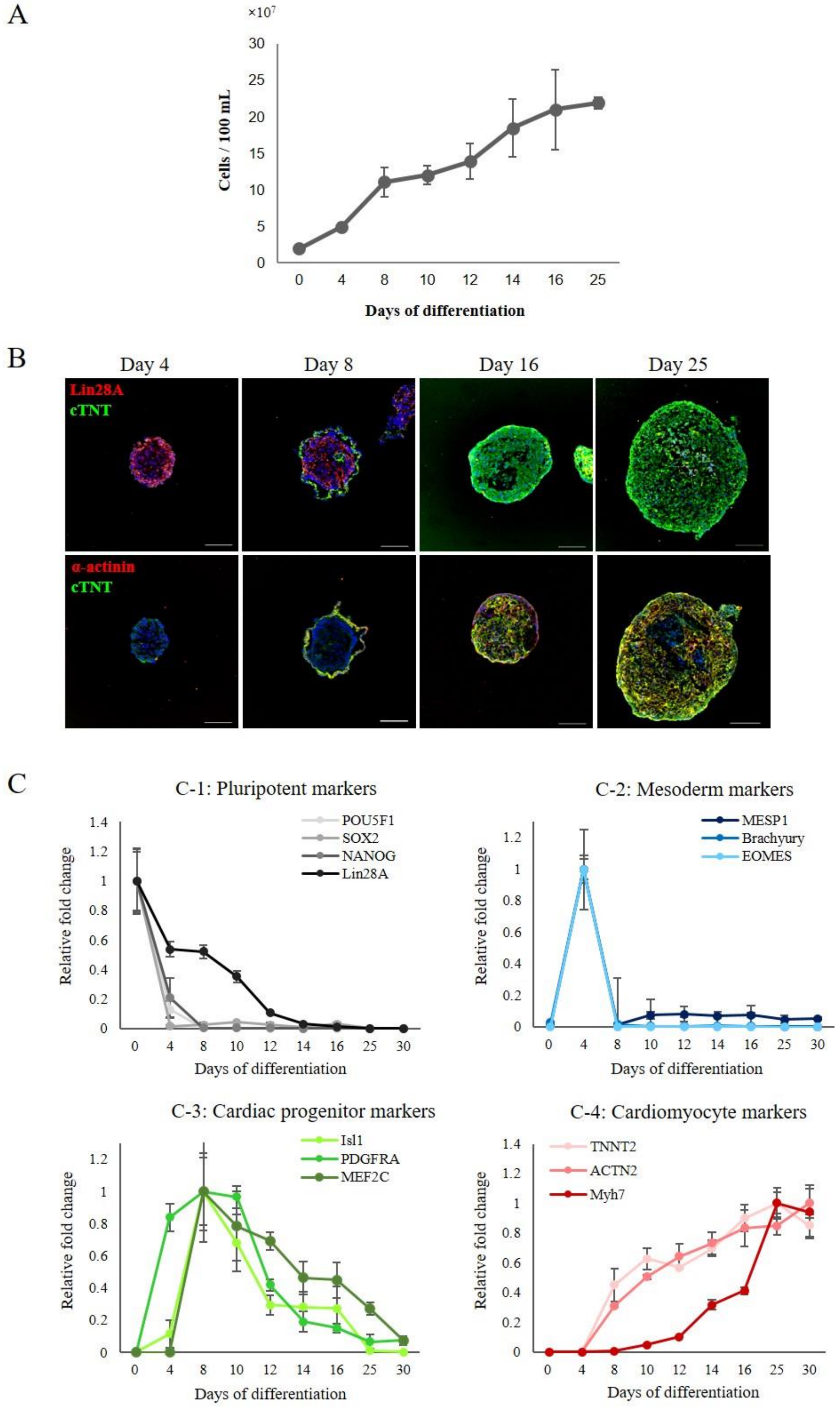

**Figure S3. Gene expression patterns of hiPSC-CM**

A

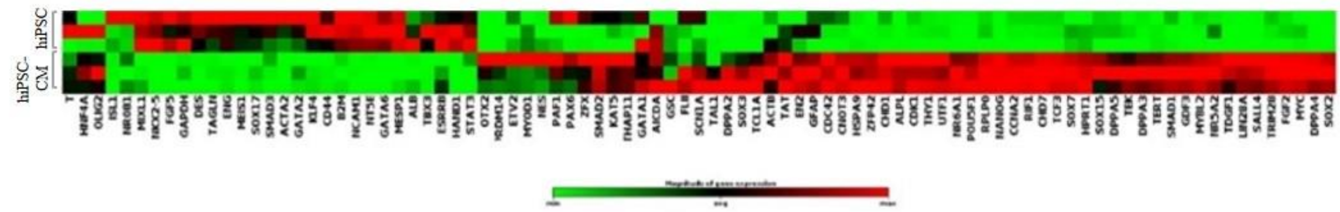


B

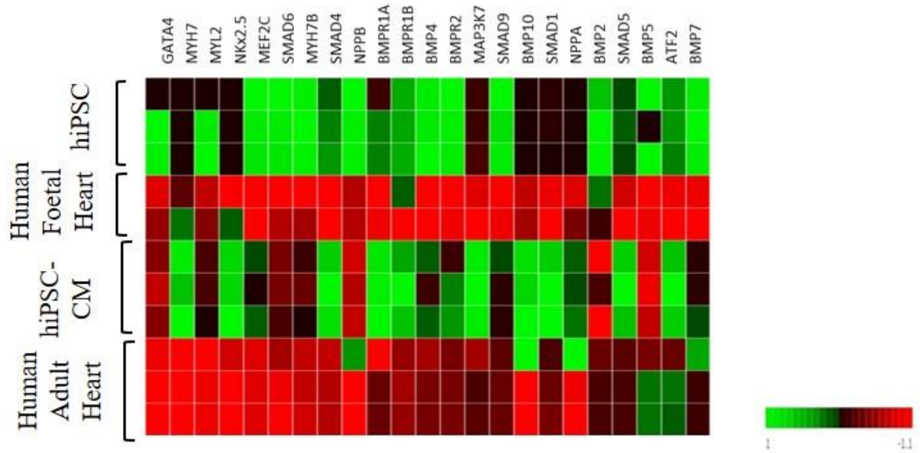


C

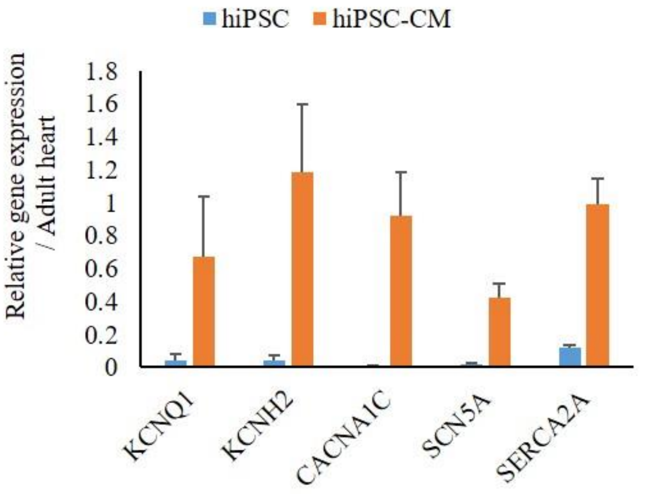

**Figure S4. Intracellular calcium levels of hiPSC-CM following drug administration**

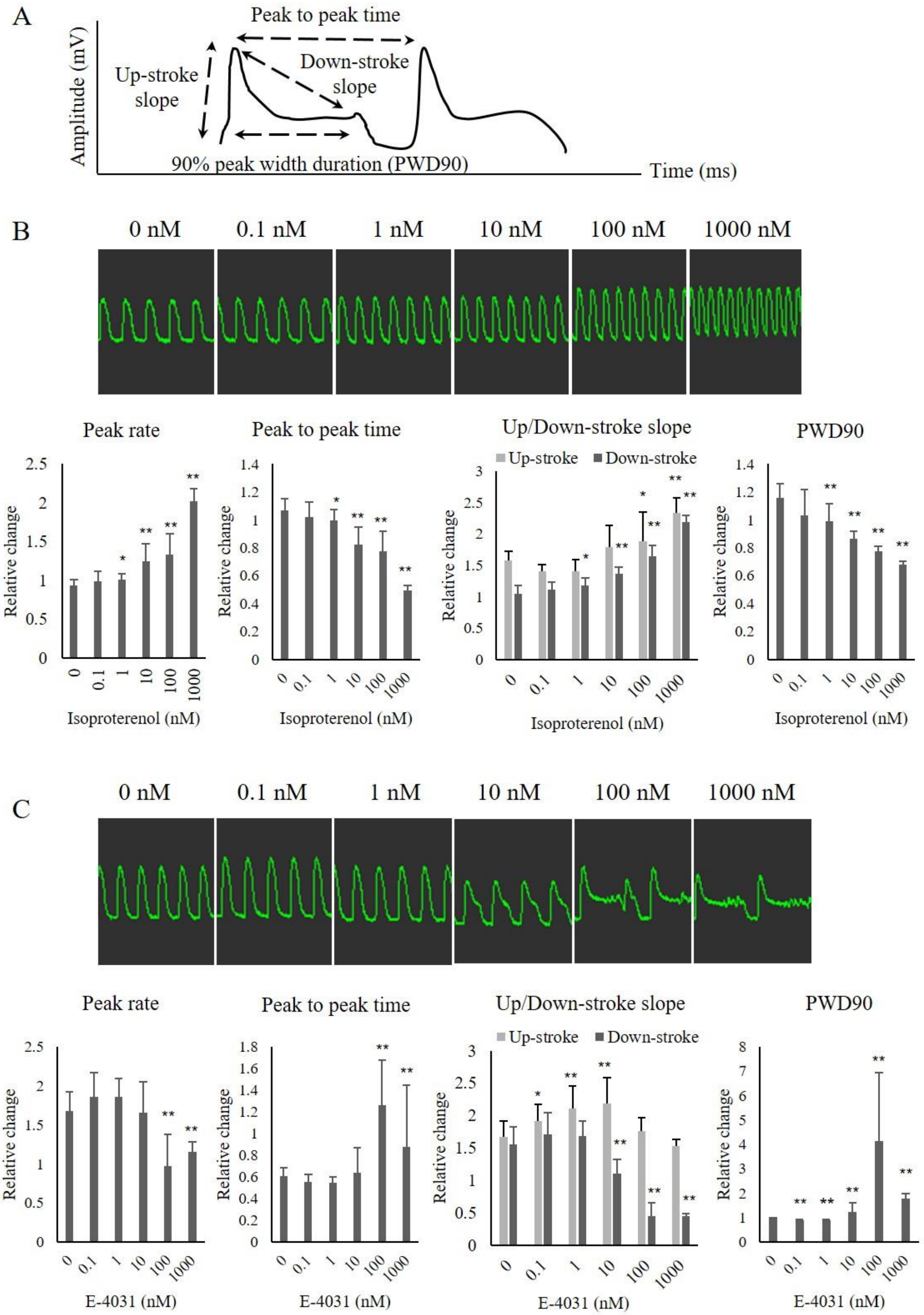

**Figure S5. Contraction properties of hiPSC-CM following drug administration**

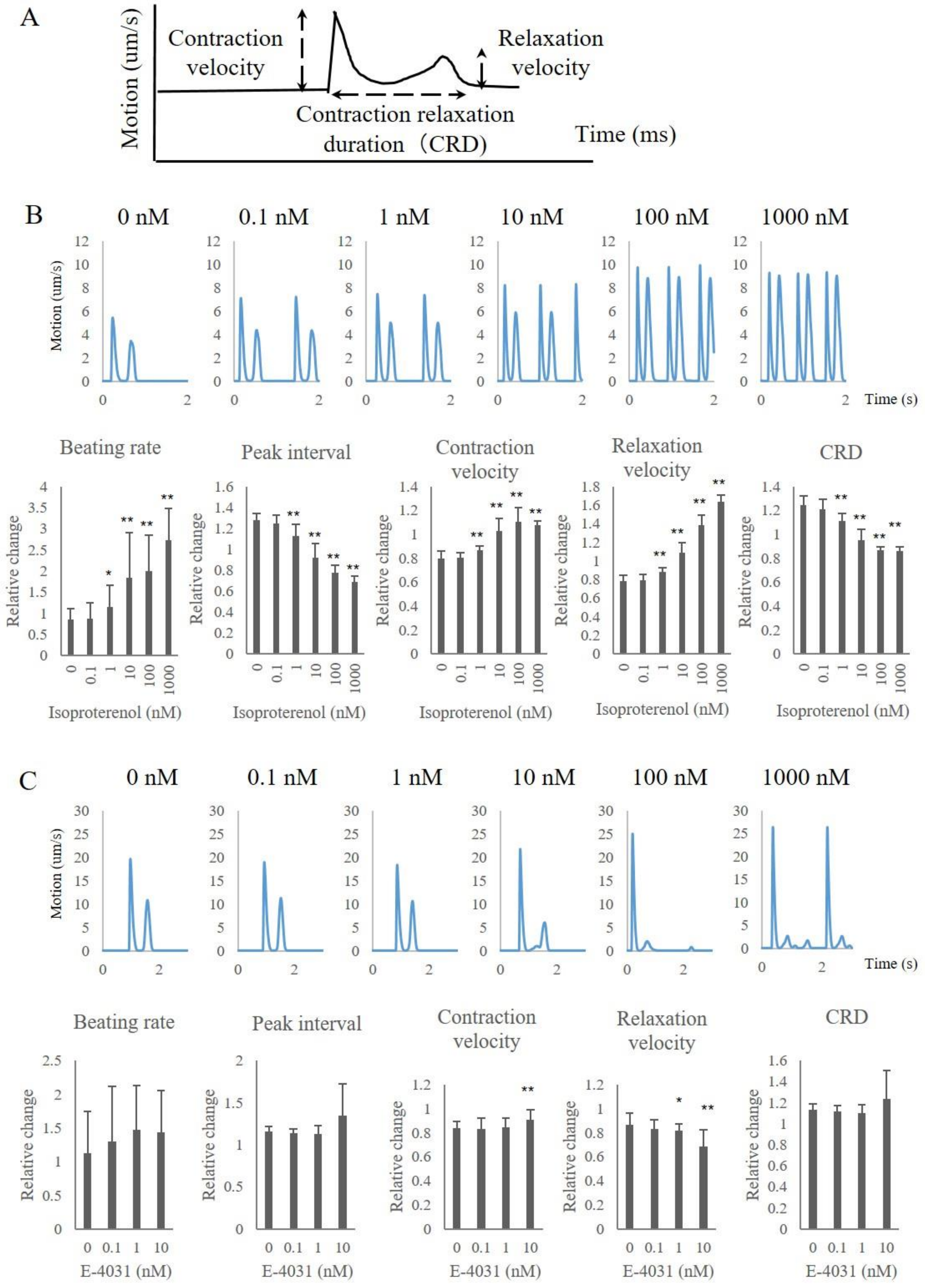

**Figure S6. Characterisation of hiPSC-CM patch**

A

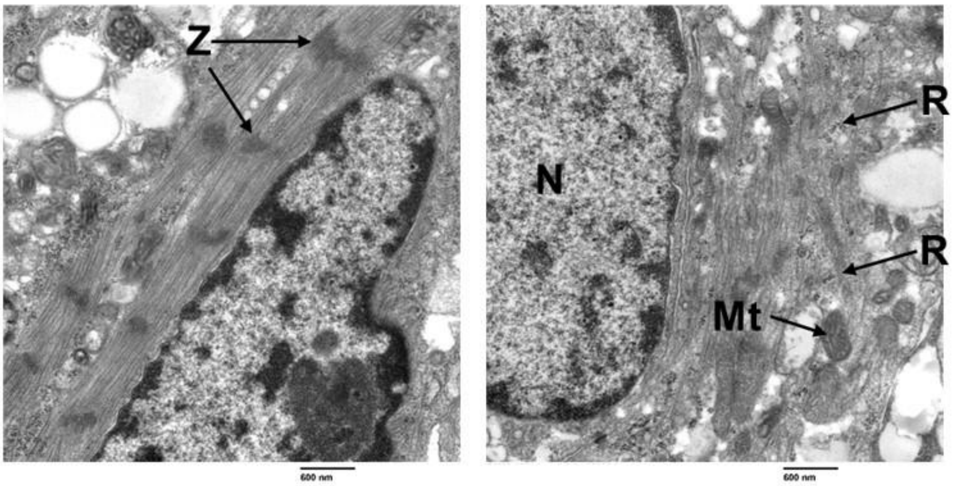


B

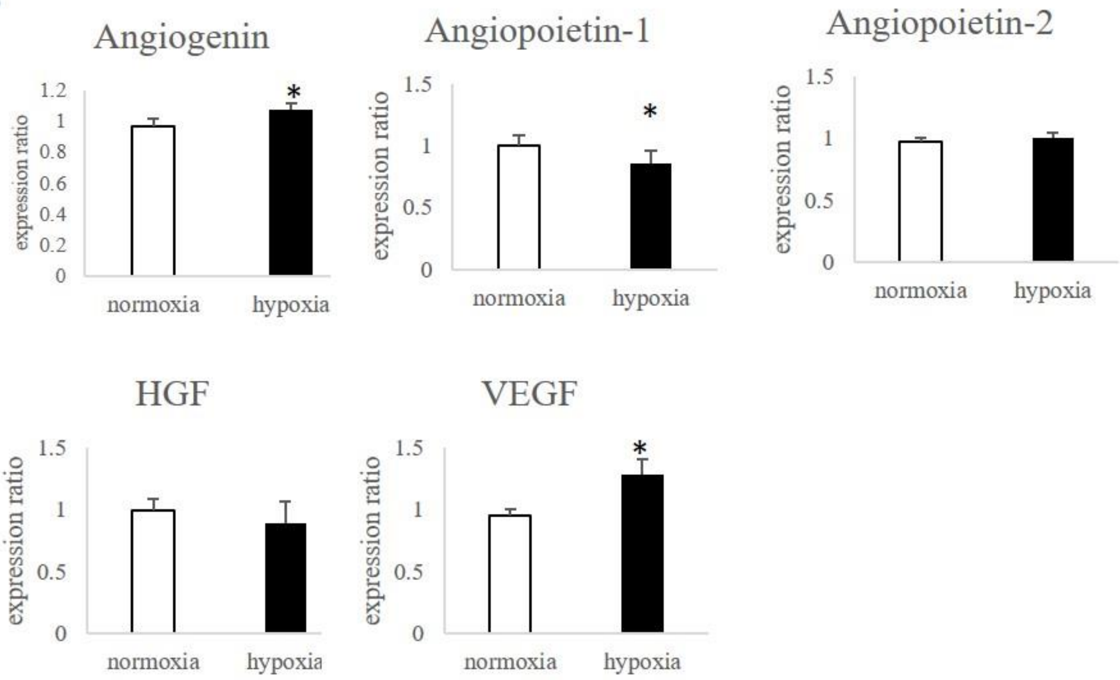

**Figure S7. Experimental protocol for determining efficacy using the porcine MI model**

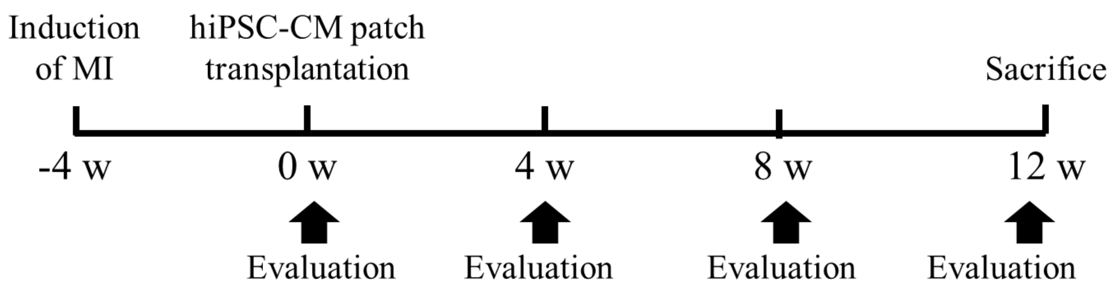


| | 0 w | 4 w | 8 w | 12 w |
|---|---|---|---|---|
| Echocardiography | ✓ | ✓ | ✓ | ✓ |
| Cardiac catheter | ✓ | | | ✓ |
| MRI | ✓ | | | ✓ |

Immunosuppressant ··············› (0 w to 12 w)

| | |
|---|---|
| Tacrolimus | 0.75 mg/kg/day |
| MMF | 500 mg/day |
| Prednisolone | 20 mg/day |

**Figure S8. Post-transplant hiPSC-CM status and effects on organs**

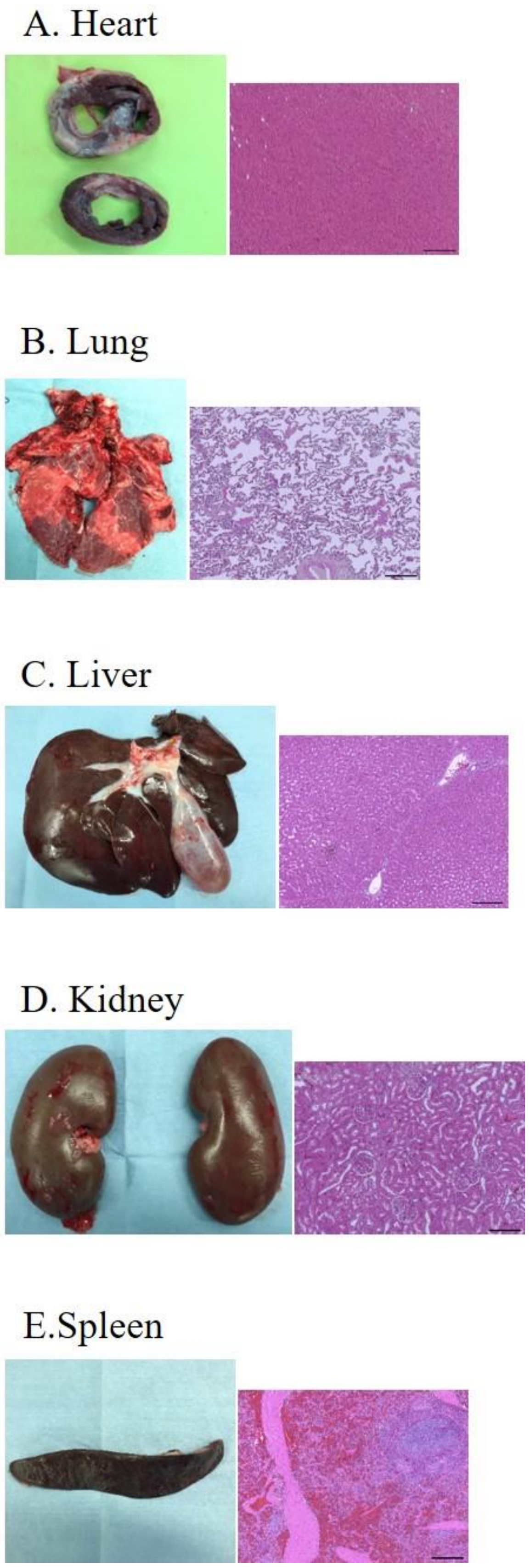
